# Tunneling nanotubes mediate mitochondrial homeostasis in cancer

**DOI:** 10.64898/2026.06.12.730900

**Authors:** Ines Saenz-de-Santa-Maria, Anna Pepe, Christel Brou, Jean-Yves Tinevez, Iakov Vitrenko, Thomas Cokelaer, Elizabeth Cohen-Jonathan Moyal, Roberto Weigert, Chiara Zurzolo

## Abstract

Tumor progression is driven by cancer cells’ ability to establish a cellular network through tunneling nanotube-like connections (TNTs), which enable mitochondrial exchange both within the tumor cells and with the tumor microenvironment (TME). However, the effects of mitochondrial transfer between tumor and non-tumor cells, and its occurrence *in vivo*, remain poorly understood. In this study, we demonstrate bidirectional mitochondrial transfer: damaged mitochondria from Glioblastoma (GBM) cells trigger mitophagy in non-tumoral astrocytes (AS), while healthy mitochondria from AS enhance the metabolic activity of GBM cells. Furthermore, intravital subcellular microscopy (ISMic) in a live animal model, allowed the visualization of TNT connections with characteristics similar to those observed *in vitro* and supported TNT-mediated mitochondrial transfer *in vivo*. These findings provide critical insights into TME interactions and their contribution to cancer resilience, highlighting TNTs as a potential target for therapeutic intervention and paving the way for further research into their role in cancer.

## Introduction

Mitochondria are double membrane bound organelles crucial for cell life and death^1^. In addition to producing ATP via oxidative phosphorylation (OXPHOs), they also regulate programmed cell death and are the main source of reactive oxygen species (ROS)^2^. Mitochondrial exchanges either among tumor cells, or among tumor and non-tumoral cells in the tumor microenvironment (TME), have been implicated in diverse processes, including metabolic reprogramming, immune suppression, cancer progression, modulation of therapeutic responses, activation of cellular rescue mechanisms, and evasion of apoptosis in recipient cells^3–16^.

A primary mechanism for mitochondria horizontal transfer occurs via transient intercellular structures known as tunneling nanotubes (TNTs)^17^. TNTs, described *in vitro* in a wide variety of tumors including Glioblastoma (GBM)^18,19^ and head and neck squamous cell carcinomas (HNSCC)^20^, are heterogeneous, highly dynamic membranous bridges that facilitate direct cytoplasmic exchanges between connected cells. They are classified into two major types: (1) Actin-based TNTs, which are short (up to 100 µm), thin (≤0.7 μm), and dynamic^21^; and (2) Microtubule (MT)-based TNTs, which are longer (up to 250 µm), thicker (≥0.7 μm), and more stable^13,22^. The latter increase in cancer cells when subjected to stress and have been proposed to favor survival of cells damaged by ischemia^23^, as well as following chemo/radiotherapy^13^. Indeed, the transfer of mitochondria between tumor cells and/or between tumor cells and non-tumoral cells by TNTs has an impact on tumor progression and resistance to chemotherapy^5,7–9,11,18,24,25^. Yet, how the composition of the TNTs influences the mitochondrial exchanges between cancer cells is unclear.

GBM is the most aggressive primary brain tumor in adults with no cure^26^. In GBM, tumor aggressiveness and progression is associated with a network of two types of intercellular connections: Tumor Microtubes (TMs), and TNTs^4,18,27^. A key feature of GBM is its complex TME, which includes non-tumoral cells such as astrocytes (AS)^28,29^. Notably, mitochondrial transfer from AS to GBM stem cells (GSCs) via TMs, enhances tumorigenesis through a mechanism depending on GAP-43 (growth associated protein enriched in growth cones)^4^,. Interestingly, TNTs have been shown to coexist with TMs in 3D GBM organoids formed by GSCs, and are hypothesized to cooperate with TMs in promoting tumor radio-resistance^18^. Consistently, TNT mediated mitochondria transfer from mesenchymal stem cells (MSC) to GSCs has been shown to induce GSC metabolic reprogramming and resistance to temozolomide (TMZ)^11^. Conversely, TNTs have also been demonstrated to mediate mitochondrial transfer from GBM cell lines to immortalized non-tumoral AS, facilitating their metabolic adaptation and hypoxia response within the TME^25^. Furthermore, AS have been shown to play a neuroprotective role by degrading damaged mitochondria ^30^ and transferring healthy ones to neurons ^31^.

Based on these findings, we hypothesize that GBM cells exploit TNTs for the bidirectional transfer of both healthy and damaged mitochondria with non-malignant cells in the TME, particularly AS, as a mechanism to enhance cancer cell survival.

To date, the existence of TNTs in cancer *in vivo* has been questioned due to technological limitations that made their detection arduous in living animals. The optical resolution of classical microscopy does not allow for the morphological characterization of these thin connections in a tissue environment, and there is currently no TNT-specific marker available^22^. Only TM, thicker neurite-like connections, have been visualized in cancer^27^ *in vivo.* These structures have recently been proposed to mediate mitochondria transfer, by confocal imaging of fixed tissue samples^4^. Consequently, whether functional TNTs facilitate mitochondrial transfer in tumors in living animals remains an open and challenging question.

Here we investigated whether different types of GBM cells communicate with each other and with AS through TNTs, focusing on the horizontal transfer of mitochondria. We used various cell models, including U251 GBM cell line, patient derived GSCs: GSC (-) and GSC (+) (also known as SRC1 and SRC2), with different metabolic activities and responsible of tumor relapse^18,32^, as well as primary murine AS, to investigate bidirectional mitochondrial transfer. Our findings revealed that damaged mitochondria from GBM cells undergo mitophagy after transfer in AS. Conversely, RNA sequencing of GSCs receiving AS-derived healthy mitochondria demonstrated upregulation of mitochondrial metabolic pathways, supporting a role for AS derived-mitochondria in enhancing the metabolic functions of GBM cells.

To investigate whether mitochondrial transfer via TNTs occurs in cancer *in vivo*, we employed Intravital Subcellular Microscopy (ISMic) a cutting-edge technique that enables real-time visualization of biological processes in live animal^33–35^. Using this approach, we observed TNTs exhibiting characteristics similar to those identified *in vitro,* containing mitochondria within the structures. Furthermore, we detect exogenous mitochondria in the receiving cell population, providing direct evidence of mitochondrial transfer *in vivo*.

Our multidisciplinary approach, which combines *in vitro* and *in vivo* systems, provides significant insights into TNT-mediated interactions within TME and their role in cancer resilience, underscoring the potential of targeting this process in future therapeutic strategies.

## Results

### Dynamic analysis of TNT-mediated mitochondrial transfer between live cancer cells

TNT-mediated intercellular communication between distant cells is well documented in several tumor cells *in vitro*, including GBM^18,20,36,37^. However, most of the features of the TNTs including diameter, length, and cytoskeletal composition were determined in fixed samples. Since chemical fixation partially disrupts the thinner TNTs supported exclusively by actin filaments (here referred as Actin-based TNTs), we used time-lapse imaging to study: i) the dynamics of TNT-mediated mitochondrial transfer between different cells, and ii) the TNT cytoskeletal composition and its role in TNT-mediated cell-to-cell communication. To this end, we used U251, the most widely used GBM-derived cell line, and GSCs derived from infiltrative regions of the same GBM patient, which exhibit different metabolic activities (GSC - and GSC +) and have been linked to relapse^32^. First, we co-transfected U251 cells with an inner mitochondrial membrane marker fused to RFP (pLV-mitodsRED) and the F-actin sensor Life-Act fused to iRFP670 (pLV-lifeAct-670). We found that cells established TNT-like connections that allowed mitochondria to be transferred (Fig. 1A and Supplementary Video S1) and observed that mitochondrial speed varies throughout the transfer process, exhibiting a saltatory motion with frequent stopping phases (Fig. 1B, a-b). Mean-square displacement (MSD) measures of mitochondrial trajectories suggested that mitochondrial transfer is due to active transport through the connections (MSD slope curve > 1, Figure 1B-c)^38^. The average max speed of mitochondria was 13.45 ± 13 nm/s (N=16) consistent with previously reported mitochondrial migration speeds via TNTs^20,39^ (Fig. 1C-a). Notably, we found that individual mitochondria inside TNTs transferred significantly faster than mitochondria moving in groups (average max speed of 26.88 ± 23 nm/s, average speed of 11.53 ± 10 nm/s, Graph 1C-b). Due to the movements of the connected cells the TNT length varied over time. The average length of TNTs in U251 cells was 35.58 ± 21 µm, with a maximum of 118.9 µm and a minimum of 9.21 µm (Fig. 1D-a). Importantly, the speed of the mitochondria transfer was independent of the TNT-length (Fig. 1D-b).

**Fig. 1.**
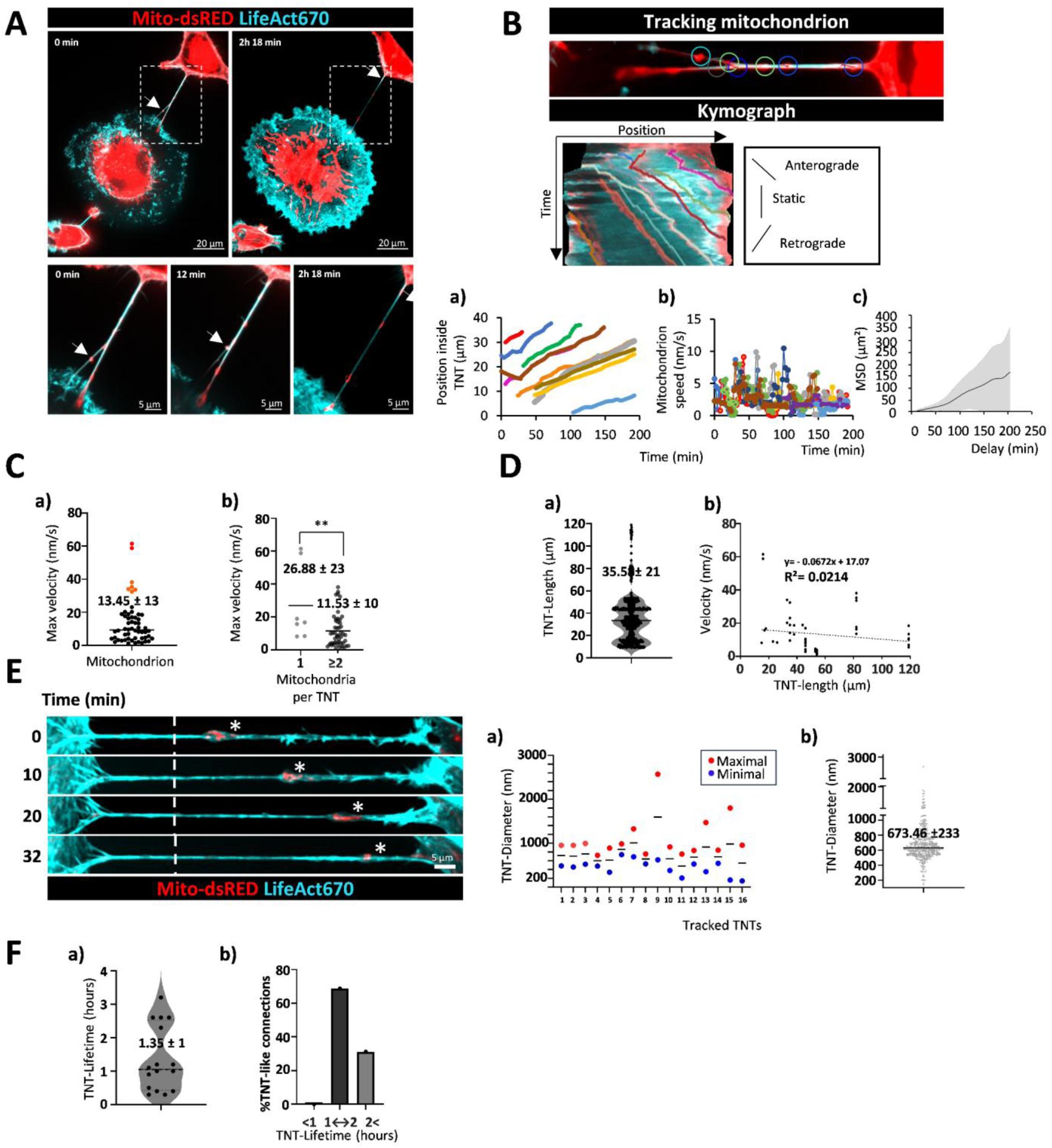
| In-depth analysis of mitochondrial transfer via TNTs in U251 cells. (A) Time-lapse images of mitochondrial transfer through TNTs in U251 cells co-transduced with pLV-lifeAct-670 (actin, light blue) and pLV-mitodsRED (mitochondria, red). Magnified images show mitochondrial movement in a branched TNT structure (see Supplementary Video S1). White arrows indicate mitochondrial migration. Scale bars: 20 µm (top), 5 µm (bottom). (B) Mitochondrial tracking and kymograph generation using the TrackMate plugin. Tracking lines in the kymograph represent different mitochondrial trajectories (anterograde, retrograde, static). Quantification of (a) speed and (b) distance traveled, and (c) MSD plot from 18 mitochondrial tracks across 9 independent experiments. The MSD slope >1 indicates active transport (One-sided t-test p-value = 1.1e-16). (C) Maximum mitochondrial speed during TNT transfer. (a) Dot plot showing the maximal speed of 56 mitochondria tracks, with an average speed of 13.45 ± 13 nm/s (red dots indicate higher speeds, with a maximum of 61.52 nm/s). (b) Speed comparison for single vs multiple mitochondria in the same TNT (**p < 0.005, unpaired t-test). (D) (a) Violin plot showing TNT length (mean = 35.57 ± 21 nm); (b) correlation between mitochondrial speed and TNT length (linear regression, R² = 0.0214). (E) TNT diameter analysis over time. The diameter was measured using the lifeAct channel intensity profile, excluding membrane bulging (see Supplementary Video S2). (Left) Representative images showing TNT diameter measurements at different time points. White dashed lines indicate measurement points, with white asterisks marking membrane bulging filled with mitochondria. Scale bar: 5 µm. (Right) (a) Maximal and minimal TNT diameters from 16 TNTs, showing that the minimum diameter is <1 µm. (b) Dot plot of TNT diameters at different time points (mean diameter = 673.46 ± 233 nm). (F) TNT persistence analysis. (a) Lifetime distribution (mean = 1.35 ± 1 hour) and (b) TNT duration categories (>1 hour, 1↔2 hours, <2 hours). Total TNTs analyzed: 16; total diameter measures: 462; total length measures: 530. N = 12 independent biological experiments.

A key feature distinguishing TNTs from other types of connections, such as TMs, is their diameter. TNT diameters ranged from 20 to 700 nm (thin canonical TNTs) and up to 1 µm (thick TNTs)^22^. In several TNTs, we observed membrane dilations (previously termed gondolas^40^) filled with mitochondria moving along the TNT during mitochondrial transfer (Fig. 1E, Supplementary Video S2). These gondolas temporarily increased the diameter of the TNT and facilitated the exchange of globular or tubular mitochondria, with diameters ranging from 0.5 to 1 µm. We measured TNT thickness over time throughout the entire movie, excluding gondolas. All connections were within the typical size range of TNTs, with an average diameter of 673.46 ± 233 nm (Fig. 1Ea-b) supporting the idea local membrane remodeling (ei., gondolas) facilitates the transfer of mitochondria through thin TNTs (<700 nm).

Several studies have shown that TNTs can persist from several minutes to several hours^20,22^, with their stability influenced by actin dynamics, presence of MT^39^ or other proteins such as EPS8, IRSp53^41^, and CD9^42^. The average TNT lifetime was 1.35 ± 1 hours, with the longest recorded lifetime reaching up to 7 hours in rare cases (Fig. 1Fa-b, Supplementary Video S3). Moreover, we confirmed that TNTs formed between two distinct cells and not between daughter cells post-division, as shown by co-culturing cells expressing lifeAct fused with different fluorophores, 670 and BFP2 (Supplementary Fig. S1, Video S3 and S4).

### TNT-transferred mitochondria integrate into the recipient cell’s mitochondrial network

Next, we sought to determine whether the cytoskeletal composition of TNTs in GB/GSC affects mitochondrial transfer. To simultaneously label mitochondria, actin filaments, and MT, we co-transduced U251, GSC (-), and GSC (+) cells with three different plasmids: pLV-mito-dsRED, pLV-lifeAct fused to either BFP2 (blue) or iRFP670 (far-red), and pLV-GFP-tubulin (Fig. 2 A). The expression of these probes did not disrupt the normal mitotic behavior of the cells (Supplementary Fig. S2, Video S5). The nature of TNTs appears to be cell specific. One-third of the TNTs in U251 and GSC (+) cells (32.76 ± 7% and 21.52 ± 10, respectively), contained both F-actin and MT (MT-based TNTs), while the remaining contained only F-actin (Actin-based TNTs). In contrast, GSC (-) contained primarily Actin-based TNTs (96.20 ± 4%) (Fig. 2B).

**Fig. 2.**
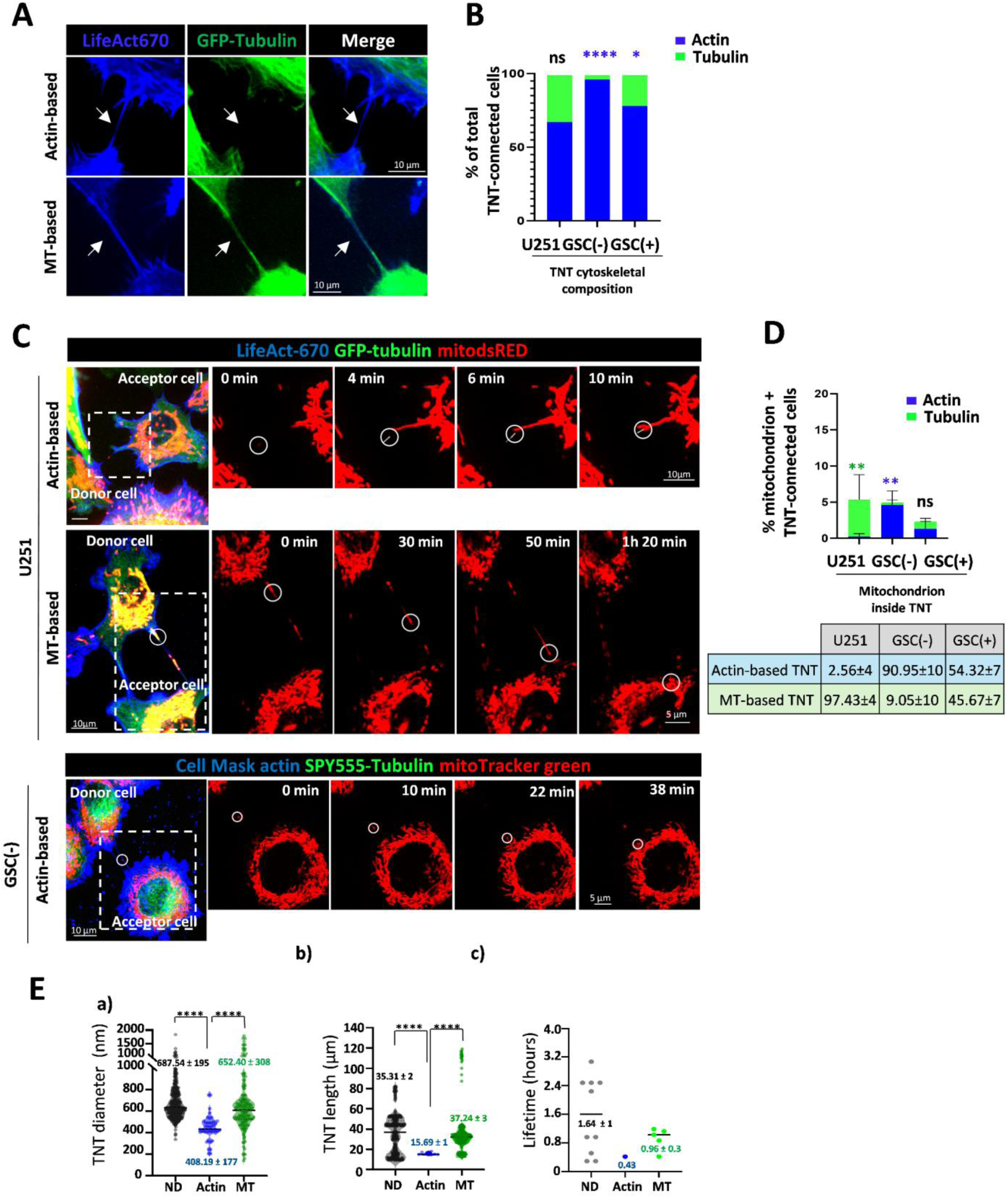
| Mitochondria transferred via TNTs fuse with the recipient cell’s mitochondrial network, regardless of TNT cytoskeletal composition. (A) Representative images of U251 cells co-transduced with pLV-lifeAct-670 (blue) and pLV-GFP-tubulin (green), showing thin Actin-based TNTs (actin only) and thicker MT-based TNTs (actin and MT). White arrowheads indicate TNTs. Scale bar, 10 µm. (B) Proportion of Actin-based (blue) and MT-based (green) TNTs in U251 and GSCs (GSC - and GSC +). GSCs predominantly form Actin-based TNTs, while no significant differences were observed in U251 cells. Two-way ANOVA test. ****p < 0.0001; *p = 0.0417; ns, p = 0.0604. Total cells counted: U251 = 807, GSC (-) = 1313, and GSC (+) = 1295. N = 3 independent experiments. (C) Time-lapse recording of mitochondria transfer via Actin-based and MT-based TNTs, illustrating their fusion into the recipient cell’s mitochondrial network. Left: Global image at time 0 shows mitochondria within an Actin- or MT-based TNT. Right: Magnified view of the red channel highlighting mitochondrial movement (white circle) over time until integration into the recipient mitochondrial network (see supplementary videos S6-S8). (D) Percentage of TNT-connected cells containing mitochondria in the indicated cell lines. Green bars: MT-based TNTs; blue bars: Actin-based TNTs. Mitochondrial transfer occurred predominantly via MT-based TNTs in U251 (green asterisk) and Actin-based TNTs in GSC (-) (blue asterisk), with no significant differences observed in GSC (+). p < 0.005 (green = 0.0043; blue = 0.0094). The bottom table shows the proportion of different TNT types containing mitochondria in the three cell types. N = 3 experiments. Two-way ANOVA. Total cells: U251 = 807, GSC (-) = 1313, GSC (+) = 1295. (E) Quantification of thickness, length, and lifetime of ND-, Actin-based, and MT-based TNTs in U251 cells by live imaging. TNT thickness was 687.54 ± 195 nm, 408.19 ± 177 nm, and 652.40 ± 308 nm; TNT length was 35.31 ± 2 µm, 15.69 ± 1 µm, and 37.24 ± 3 µm; and TNT lifetime was 1.64 ± 1 hours, 0.43 ± 0.1 hours, and 0.96 ± 0.3 hours, respectively, for ND-based, Actin-based, and MT-based TNTs. ****p < 0.00005, Two-way ANOVA, multiple comparisons. Total analyzed TNTs transferring mitochondria: 16 (ND-based = 10, Actin-based = 1, MT-based = 5). N = 12 independent biological experiments.

Both types of TNTs supported the transfer of mitochondria from donor (D) to recipient cells (Fig. 2C, Supplementary Fig. S3, and Videos S6-S7 and S8 for Actin- and MT-based TNTs, respectively), although the extent of the transfer was cell-type dependent. Specifically, in U251 cells we found that MT-based TNTs transported the vast majority of the mitochondria present in TNTs (97.43 ± 4%), despite the higher frequency of Actin-based TNTs cells (Fig. 2D). On the other hand, in GSC (-) mitochondria were transported almost exclusively by Actin-based TNTs (90.95 ± 10%), whereas in GSC (+) both kind of TNTs equally contributed to the mitochondrial transport (45.67 ± 7% MT-based, 54.32 ± 7% Actin-based). Notably, these results show that the transport of mitochondria does not correlate with the number of a particular kind of TNTs.

We also analyzed the diameter, length, and lifetimes of the different mitochondria-transferring TNTs. Our results showed that TNTs supported by MT exhibited significantly increased diameter and length compared to Actin-based TNTs, with no significant difference compared to non-determined (ND-TNTs), formed between cells expressing mito-dsRED-lifeAct670 (labelling mitochondria and actin filament) but not GFP-tubulin.

The average diameter and length for Actin-based TNTs were 408.19 ± 177 nm and 15.69 ± 1µm, respectively; for MT-based TNTs: 652.40 ± 308 nm and 37.24 ± 25 µm; and for ND-TNTs 687.54 ± 195 nm and 35.31 ± 19 µm (Fig. 2E). The lack of significant differences between MT- and ND-TNTs was consistent with the results of Figure 2D showing that mitochondrial transfer in U251 cells occurs predominantly via MT-based TNTs and suggesting that most ND-TNTs are actually MT-based TNTs.

Finally, we found that ND- and MT- populations of TNTs had an average lifetime of 1.64 ± 1 hours and 0.96 ± 0.3 hours, respectively, whereas Actin-based TNTs only 26 min (Fig. 2E), supporting the hypothesis that MTs increase the stability of TNTs^22^. In all cases, no matter whether transfer occurred through MT-based or Actin-based TNTs, transferred mitochondria successfully integrated the mitochondrial network of the recipient cell (Fig. 2C, Supplementary Videos S9 and S10 for MT- and Actin-based TNTs, respectively).

### Mitochondria transfer bidirectionally between tumor and healthy cells through TNTs

To assess whether tumoral cells communicate via TNTs with non-malignant cells from the TME, we decided to focus on AS because they make up 40-50% of the glial cells in the brain and play a crucial role in supporting neurons ^31^. Moreover, Neuzil’s group have recently identified AS and not microglia as the predominant mitochondrial donors in the brain^15^. We co-cultured non-tumoral primary murine AS (Supplementary Fig. S4A, B) with GB/GS cells. We observed that both homotypic (GB/GS-GB/GS or AS-AS) and heterotypic (GB/GS-AS) TNTs were established in 2D-culture and in tumor organoids (Fig. 3A, Fig S4 C-E). Moreover, individual tumor cells formed heterotypic and homotypic connections simultaneously (Fig. 3A). To determine the directionality of the transfer, we transfected one of the cell populations (termed D) with a GFP-tagged mitochondrial marker (mito-GFP).

**Fig. 3.**
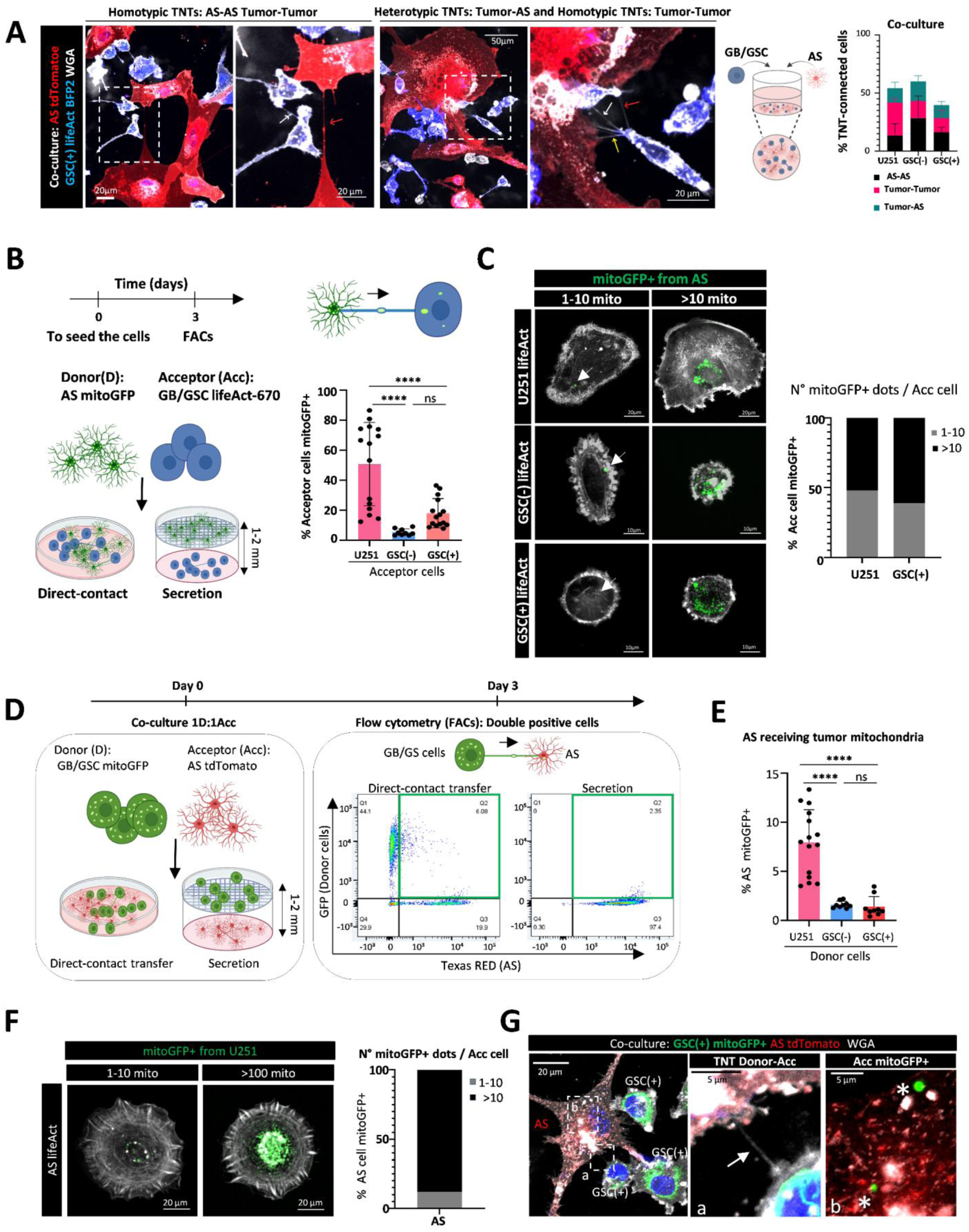
| GB/GS cells form functional TNT-heterotypic connections with primary cortical AS, facilitating mitochondrial transfer. (A) Representative images of co-cultures of GSC (+) cells expressing pLV-lifeAct-BFP2 (blue, actin filaments) and primary cortical AS tdTomato (red), as shown in the schematic. Cells were plated overnight, fixed, and stained with WGA (white) to label cell membranes. GSC (+) and AS form both homotypic TNTs (linking either tumor cells (GSC+, Tumor-Tumor) or non-tumor cells (AS-AS)) and heterotypic TNTs (Tumor-AS). Magnified views of the white-dotted squares highlight homotypic TNT connections (white and red arrowheads) and heterotypic TNTs (yellow arrowheads). Scale bars: 20 µm (left), 50 µm (right), and 20 µm (magnifications). The graph shows the percentage of TNT-connected cells in co-cultures of AS with U251, GSC (-), and GSC (+), including homotypic TNT (AS-AS in black, Tumor-Tumor in pink) and heterotypic TNT (Tumor-AS in green). Representative images of U251 and GSC (-) co-cultures are in Supplementary Fig. S4D. (B) Experimental timeline for analyzing mitochondrial transfer from AS to tumor cells (U251, GSC (-), and GSC (+)). AS expressing mitoGFP were co-cultured with tumor cells expressing LifeAct at 1:1 ratio, either in direct contact or separated by a 1 µm filter (secretion control). After three days, FACS was used to quantify double-positive cells, representing tumor cells that internalized astrocytic mitochondria. On the right, FACS quantification shows the percentage of tumor cells receiving mitochondria via direct contact, after subtracting transfer due to secretion. Results show 50.72 ± 28% of U251 cells received mitochondria from AS, significantly more than GSC (-) (5.43 ± 2%) and GSC (+) (17.99 ± 10%) cells. p < 0.0001, ns = 0.0959 (two-way ANOVA, multiple comparisons). (C) Representative images of tumor cells containing astrocytic mitochondria immediately after sorting. Left: Tumor cells with 1-10 mitochondrial particles vs. cells with >10 particles per cell. White arrows indicate cells with a single particle. Scale bars: 20 µm (U251) and 10 µm (GSC). Right panel: Quantification of mitochondrial particles per tumor cell in U251 and GSC (+), showing cells with 1-10 mitochondria (gray) and >10 (black). (D) Experimental timeline for analyzing mitochondrial transfer from tumor cells (U251, GSC (-), and GSC (+)) to AS. Experiments were performed as in (B), but with tumor cells and AS as donor and acceptor cells, respectively. Representative FACS plots show results for direct contact co-cultures (left) and secretion controls (right). Acc and D cells are plotted on the X and Y axes, respectively, with Acc cells containing internalized mitochondria gated in green. (E) FACS quantification of mitochondrial transfer from tumor cells (D) to AS (Acc). Results show 7.95 ± 3% of AS received mitochondria from U251 cells, significantly more than from GSC (-) (1.6 ± 0.3%) and GSC (+) (1.39 ± 1%). p < 0.0001, ns = 0.9712 (two-way ANOVA, multiple comparisons). (F) Representative image of AS containing tumor-derived mitoGFP+ particles immediately after sorting. The graph shows the percentage of AS with 1-10 mitochondria (gray) and >10 mitochondria (black). Scale bar: 20 µm. (G) Co-culture of GSC (+) mitoGFP (green) and AS tdTomato (red) after three days. Cells were fixed, stained with WGA (white), and imaged by confocal microscopy 60x magnification. Inset (a) shows a TNT (white arrow) connecting a GSC (+) cell to an AS cell. Inset (b) highlights an AS cell containing tumor-derived mitochondria (AS mitoGFP+, white asterisks). Scale bars: 20 µm (left) and 5 µm (insets).

Since mitochondria can be transferred by extracellular vesicles, we used a co-culture system where both populations were separated by a 1 µm diameter filter, but in contact with the same culture medium, as a secretion control. This filter prevents direct contact between D and acceptors (Acc) cells while allowing the passage of mitochondria (0.5 -1 µm diameter). The percentage of mitochondrial transfer in both conditions—secretion and normal co-culture—was assessed after 3 days using flow cytometry (FACS) by quantifying the double-positive cells. The final percentage of Acc cells receiving mitochondria by direct contact was calculated by subtracting the percentage of mitochondrial transfer via secretion from the overall percentage (Fig. 3B).

We found that AS, when examined as D, transferred mitochondria to 50.72 ± 28% of the U251 cells, as well as 5.43 ± 2% and 17.99 ± 10% to GSC (-) and GSC (+), respectively (Fig. 3B). The transfer was also confirmed by microscopy (Fig. 3C). Conversely, when AS were examined as Acc, the transfer from GB/GS cells was less efficient. Indeed, only 7.95 ± 3% of the AS cells received mitochondria from U251 cells, and in the case of GSC (-) and GSC (+), 1.60 ± 0.3 and 1.39 ± 1%, respectively (Fig. 3E). Also, in this case we confirmed the transfer of tumor derived-mitochondria by confocal microscopy (Fig. 3F, G).

These data show that mitochondria can be transferred bidirectionally between AS and tumor cells, with the transfer from AS to tumor cells being quantitatively predominant.

### AS degrade dysfunctional mitochondria transferred from tumor cells via TNTs

We hypothesized that mitochondrial transfer could enable cancer cells to recruit functional mitochondria from AS and/or dispose of damaged ones. To test this hypothesis, we assessed the status of the transferred mitochondria and investigated their fate in the recipient cell. To monitor the accumulation of mitochondria over time, we extended the co-culture of mitoGFP-U251 and LifeAct670-AS to 4 days (See schematics of Fig. 4A). Then, we sorted the AS containing U251 derived-mitoGFP+ and quantified the number of mitochondria per Acc cell. The collected cells were plated for an additional 18 hours, after which mitochondrial functionality was assessed by labeling them with mitoTracker RED CMXRos or tetramethylrhodamine methyl ester (TMRM), which are retained by energetically active mitochondria^43,44^.

**Fig. 4.**
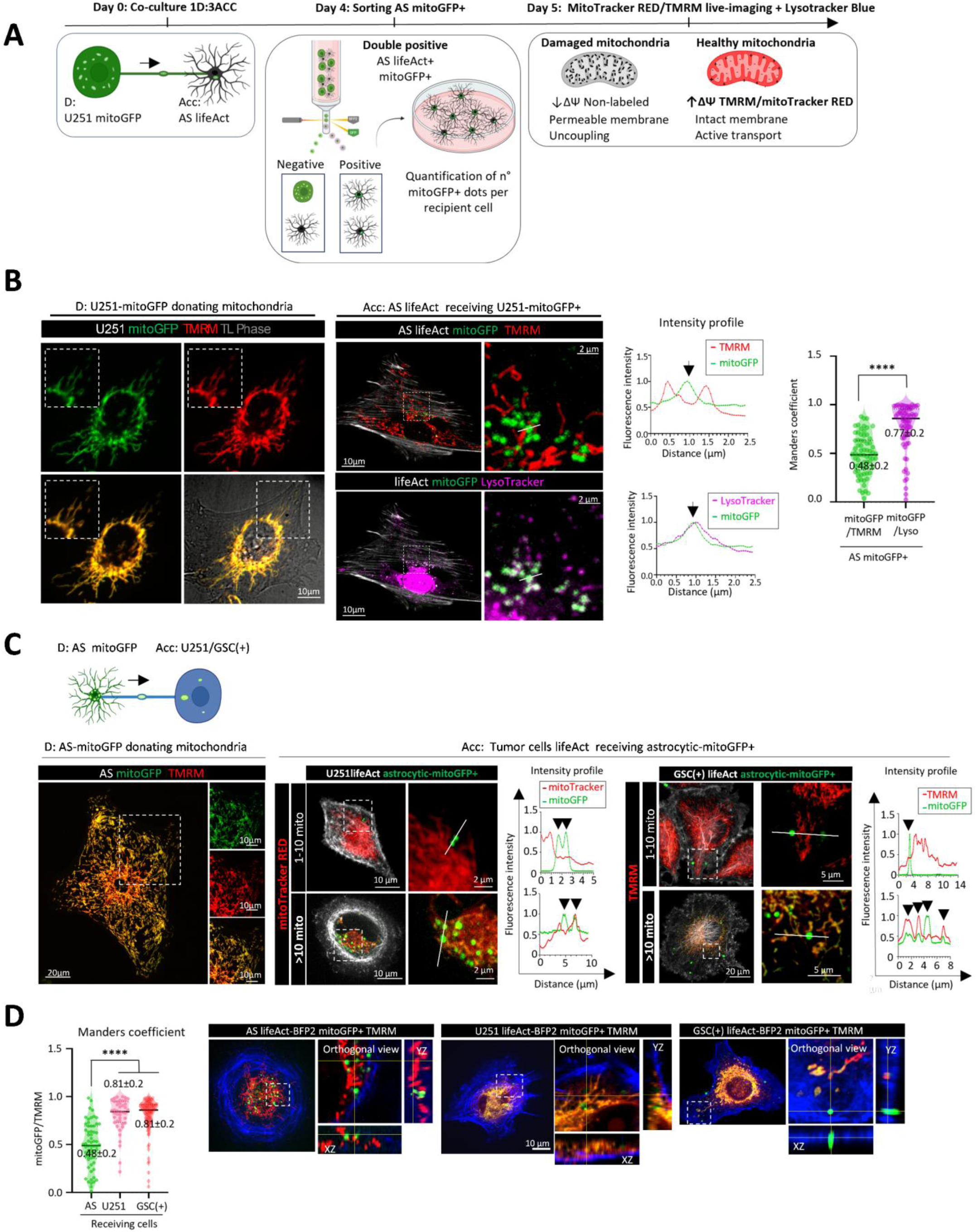
| State and fate of transferred mitochondria between AS and GB/GS cells in the recipient. (A) Experimental setup to assess the state of mitochondria transferred via TNTs from U251 cells to AS. U251 mitoGFP (mitochondria in green) were co-cultured with AS lifeAct-670 (actin filaments in far red) at 1:3 ratio. After 4 days, U251+AS cells were detached, and AS receiving U251-derived mitochondria (double-positive cells, AS lifeAct+ mitoGFP+) were sorted and seeded overnight. The next day, AS mitoGFP+ were co-stained with TMRM or MitoTracker Red CMXRos (both at 50nM, in red), which label mitochondria based on membrane potential (ΔΨ), and Lysotracker Blue DND-22 (50nM, in magenta) to track astrocytic lysosomes. Mitochondria with disrupted ΔΨ are not stained with TMRM. Live imaging analysis was performed. Scheme created with BioRender. (B) Representative images of D population U251 mitoGFP tumor cells (left) and Acc population AS (right) showing co-localization assays to assess mitochondrial state. Left: U251 cells (mitoGFP, green) show perfect co-localization with TMRM (red), forming a yellow mitochondrial network. Magnified views highlight mitochondrial details. Right: AS receiving U251-mitochondria (mitoGFP+) display triple co-localization of mitoGFP (green), TMRM (red), and Lysotracker (magenta), showing no co-localization of mitoGFP with TMRM but strong co-localization with Lysotracker, suggesting mitophagy of U251-derived mitochondria in AS. Tubular astrocytic mitochondria contrast with globular U251-derived mitochondria, indicating dysfunction. Manders’ coefficients reveal significant lysosomal association with U251-derived mitoGFP (0.77 ± 0.2) and poor TMRM association (0.48 ± 0.2). Scale bar: 10 µm. N = 3 independent biological experiments, 69 cells analyzed. Mann-Whitney test, ****p < 0.0001. (C) Representative images of AS mitoGFP (left) and tumor cells (U251, GSC (+) expressing lifeAct-670) (right) showing mitochondrial transfer state. Left: D population AS mitoGFP (green) show perfect co-localization with TMRM (red), forming a yellow mitochondrial network, with higher TMRM intensity near nuclei, indicating increased ΔΨ. Right: U251 and GSC (+) cells receiving astrocytic mitochondria (mitoGFP+). Intensity profile analysis of mitoGFP and MitoTracker Red CMXRos or TMRM in U251 and GSC (+) cells is shown. Cells receiving 1-10 or more than 10 mitoGFP particles per cell are shown. Magnified views show red and green channels for better co-localization visualization. Singular 1-2 globular mitochondria do not co-localize with ΔΨ membrane probes, while progressive integration into the mitochondrial network is seen in cells receiving more than 10 mitoGFP particles per cell. Black arrows indicate green intensity peaks. Scale bars: 10 and 2 µm (U251); 20 and 5 µm (GSC +). (D) Mitochondrial activity in recipient cells (AS, U251, GSC +) analyzed using Manders’ co-localization coefficients (mitoGFP overlapping with TMRM). Dot plot compares mitochondrial activity across cell types. Manders coefficients: 0.48 ± 0.2 (AS), 0.81 ± 0.2 (U251), 0.81 ± 0.2 (GSC +). N = 3 independent biological experiments, 62 (AS), 43 (U251), and 101 (GSC (+)) cells analyzed. Statistical significance: ****p < 0.0001, ns = 0.98 (two-way ANOVA, multiple comparisons). Right: Z-projection images of TMRM-labeled cells show mitoGFP+ dots inside recipient cells. In AS, mitoGFP+ dots remain globular and lack TMRM co-localization, suggesting inactivity. In U251 and GSC (+) cells, orthogonal views (xz and yz planes) show mitoGFP+ in green (disrupted mitochondria) and mitoGFP+ co-localizing with TMRM in yellow (active mitochondria integrated into the mitochondrial network inside recipient cells).

Colocalization analysis (Mandersˈ overlap coefficient) showed that mitochondria transferred from the GB/GS cells (mitoGFP-labelled) to the AS did not fuse with the Acc mitochondrial network, did not label with TMRM (mitoGFP/TMRM coefficient= 0.48±0.2) and were globular in shape (Fig. 4B). Moreover, they showed strong colocalization with the lysosomal marker LysoTracker Blue as shown by analysis of the Mender coefficient (mitoGFP/LysoTracker coefficient = 0.77 ± 0.2), and the fluorescence intensity profile (Fig 4B). This indicates a significant association of transferred mitochondria with lysosomes suggesting they underwent mitophagy (Fig. 4B). As expected, the native mitochondria were labelled with TMRM in both D and Acc and exhibited a tubular shape (Fig. 4B). On the other hand, mitochondria transferred from the AS into the GB/GC cells were more heterogenous (Fig. 4C). In cells where few mitochondria were transferred (<10/cells), they did not fuse with the existing mitochondria network, whereas in cells receiving a larger number of mitochondria, the transferred organelles fused with the Acc mitochondrial network, colocalized with TMRM, and maintained a tubular morphology (Fig.4 C).

These findings suggested that TNTs serve potentially to maintain viable mitochondria homeostasis in cancer cells by facilitating both the disposal of damaged mitochondria from the cancer cells and the addition of new ones from the healthy cells. This conclusion is supported by Mandersˈ colocalization analysis across the three populations studied, AS, U251, and GSC (+) (Fig. 4D).

### Astrocytic mitochondria enhance the expression of mitochondrial respiration genes in GSC

We hypothesized that TNTs help cancer cells to maintain or boost their metabolism by transferring whole healthy organelles from AS to tumor cells. To investigate this, we explored the transcriptional changes induced by TNT-dependent transfer from AS to GSC (+). We focused on GSC (+) due to their crucial role in relapse and their reliance on mitochondrial ATP synthesis^32^. We performed bulk RNA sequencing (RNA-seq) on sorted GSC (+) positive for astrocytic-derived mitochondria (mito-dsRED+) and on GSC (+) negative (mito-dsRED-) as controls (see experimental schematic in Fig. 5A).

**Fig. 5.**
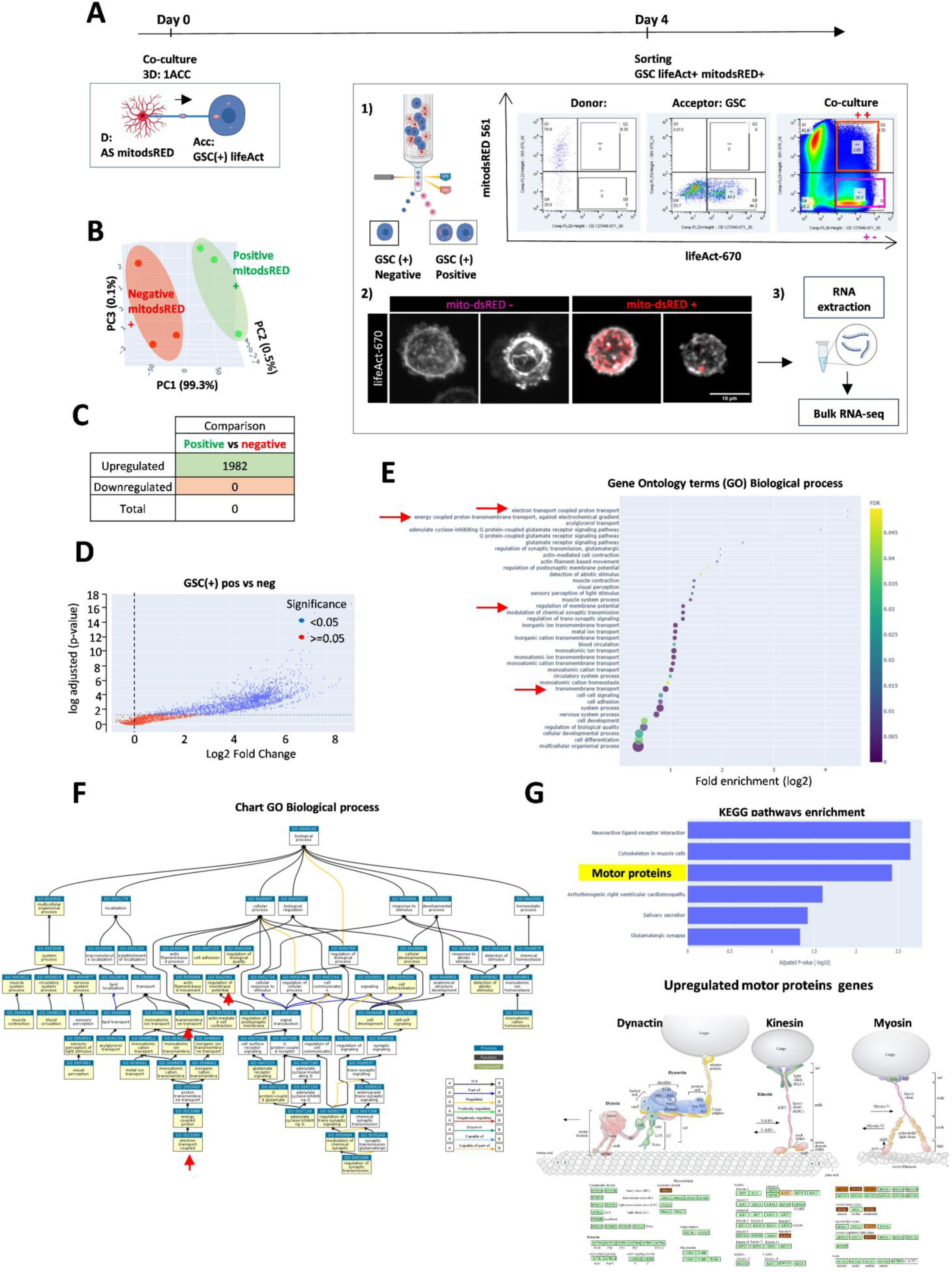
| Bulk RNA-seq analysis of sorted GSCs (+) that received mitochondria (mitoDsRED+) versus those that did not (mitoDsRED-) from AS. (A) Schematic of the experimental design of co-culture experiments and sorting of GSCs (+) with or without astrocytic mitochondria (left). Representative FACS plot of the sorting experiment and representative images of the cells (right). N= 3 independent biological co-culture experiments were analyzed. (B-F) Differential gene expression analysis (DGE). (B) PCA 3D plot of the separate samples from the different biological conditions indicating two different clusters: GSCs non-receiving mitochondria mitodsRED- (neg, red dots) and GSC recipients of mitochondria mitodsRED+ (pos, green dots). (C) Table summarizing the differential gene expression between the two clusters. (D) Volcano plot representing the differential gene expression signature of GSC astrocytic mitodsRED+ pos versus GSC astrocytic mitodsRED- neg. Blue dots indicate p-value < 0.05. Dashed lines mark fold change > 1.5 (E-F) Gene Ontology (GO) analysis of biological process (positive enrichment) for GSC pos (mitodsRED+) vs GSC neg (mitodsRED-). (E) List of principal biological process enhanced. (F) Chart GO biological process upregulated in GSC pos. Biological processes related with mitochondria metabolism are upregulated and indicated on a yellow background. (G) KEGG analysis showing 6 pathways enrichment, including motor proteins. On the right, graphic chart of motor proteins genes. Dark blue corresponds to the upregulated genes with a log2 fold change of 4 or 40.

Differential gene expression (DGE) analysis using principal component analysis (PCA) identified two clusters: control GSC (+) with no exogenous mitochondria (mito-dsRED-, neg), and GSC (+) that received astrocytic mitochondria (mito-dsRED+, pos) (Fig. 5B). These results strongly suggest that AS-GSC communication may alter the transcriptional program of GSCs.

We observed hundreds of differentially expressed genes between pos and neg clusters. Interestingly, there were 1982 upregulated genes and no downregulated genes (absolute log2 fold change above 1), indicating that the acquisition of astrocytic material acts as an activator rather than a suppressor in GSC (+) (Fig. 5C, and Volcano plot in 5D).

Gene Ontology (GO) enrichment analysis revealed the enrichment of genes related to mitochondrial metabolism, including electron transport coupled proton transport, regulation of membrane potential and transmembrane transport (Fig. 5 E, F).

KEGG pathways analysis showed that motor proteins associated with mitochondrial transport, specifically KIF6 and KIF9 (MT-based), as well as Myo3 (Actin-based), were upregulated (Fig. 5G). Additionally, axonal dynein DNAH, which is involved in the movement of cilia and flagella and retrograde movement of axon, was also identified.

These results indicate that GSC (+) cells receiving astrocytic mitochondria exhibit enhanced mitochondrial biogenesis and activity, potentially impacting their cell adhesion and motility.

### Evidence of TNT-like connections in cancer cells *in vivo* supporting their role in mitochondria exchange

TNT have been widely studied *in vitro* in several tumoral cells, in 2D- and 3D-cultures, including organoids^18^, *ex vixo* explants^45^, and tissues^46^. Visualizing the dynamics and function of these structures in live animals has been challenging. However, the development of ISMic, which offers high-resolution imaging comparable to those achieved *in vitro* provide now with this unique opportunity^33,34,47^.

Nonetheless, ISMic in live rodents—with the resolution required to detect TNTs and mitochondrial transfer—does not work in deep brain regions. To address this limitation, we used an orthotopic tongue cancer model for HNSCC suitable for ISMic^35,48^, as a tool to visualize TNTs *in vivo*.

First, we characterized TNTs in HN12 cells, a non-invasive HNSCC cell line. We found that HN12 cells form TNTs *in vitro* with characteristic such as percentage of TNT-connected cells (39.91 ± 13%), average TNT diameter (607.08 ± 227 nm), and TNT length (from 5.31 to 76.41 µm) very similar to U251 cells. Additionally, HN12 TNTs hovered above the dish, as shown in a 3D-rendering movie (Supplementary Fig. S6, Video S11).

Next, to characterize the presence and functionality (mitochondrial transfer) of TNTs in HNSCC *in vivo*. We injected 500,000 cells of a 1:1 mixture of a D population (HN12 with labeled actin and mitochondria, mitoGFP-lifeAct-BFP2) and an Acc population (HN12 with labeled actin by F-tractin mCherry) into the ventral side of the tongue of immunocompromised mice (Fig. 6A). After tumor development, we performed ISMic on the living animal at different time points up to 40 days post-injection.

**Fig. 6.**
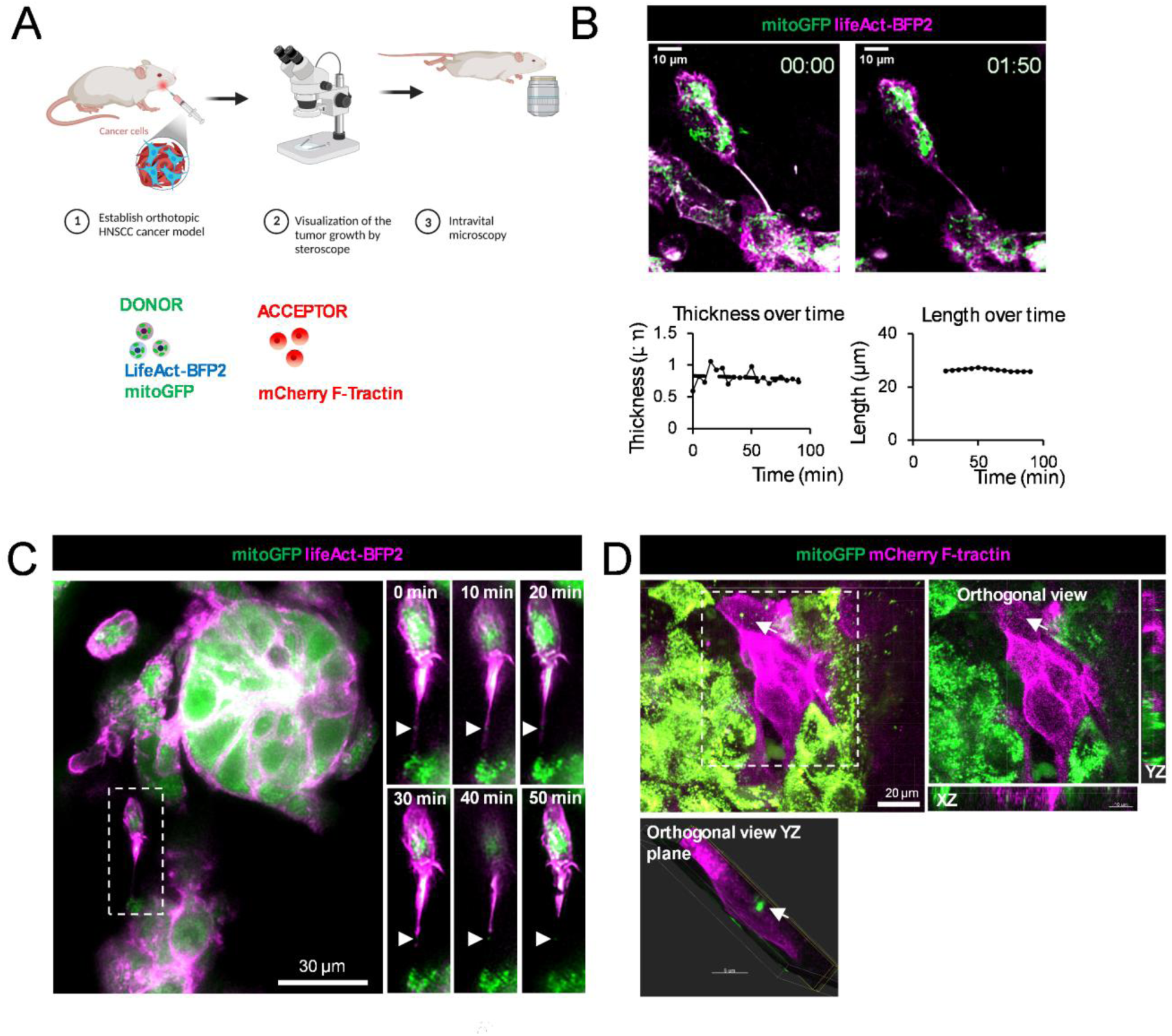
| TNT-like connection between HNSCC *in vivo.* (A) Experimental diagram of the orthotopic xenograft model used, in which (1) a mixture of engineered HN12, human-derived HNSCC cell line, was submucosally injected in immunocompromised athymic nu/nu mice into the ventral anterior side of the tongue: a D population which co-express mitoGFP and lifeAct-BFP2 (mitochondria in green, and actin filaments in blue, respectively) and an Acc population expressing mCherry F-tractin (actin filaments in red) in a 1:1 ratio. Final concentration 500,000 cells/20µl; (2) visualization of the tumor growth; (3) acquisition of time-lapse movies in the tongue of an anesthetized mouse by ISMic. (B) Representative images of a time-lapse movie at the initial and final time points show the presence of a TNT-like connection and its lifetime (upper panel) (Supplementary video S15). The quantification of the TNT variation in thickness and length over time is shown in the bottom panel. The analysis was made using TrackMate FIJI plugin. The movie was taken by two-photon microscopy, Z-stacks were acquired by a single excitation wavelength 750 nm, allowing the visualization, at the same time, of the actin filaments in emission filter 405 nm and the mitochondria in green using 505—560 filters. The movie is a 30.8 µm Z-projection of a total Z-stack of 44 with a Z-stack interval of 0.7 µm. The time interval is 5 min (total time stacks 18). Magnification 40X. Scale bar 10 µm. (C) Global image of the tumoral niches present in the animal tongue showing two cells connected via TNT with one mitochondrion inside. The right panel shows a magnification of the white square to better visualization of the TNT. Representative snapshots at the indicated time points. 1 hour of movie duration was not enough to visualize the complete transfer of mitochondria from one cell to another. White arrows point the position of the mitochondrion over time. Scale bar 30 µm. Timeframes result from the max projection of 64 slides (step size 0.7 µm) with a total physical thickness of 44.8 µm, with 4 min interval time total duration 50 min (total time stacks 12). Magnification 30X. Scale bar 30 µm. (D) Single focal plane (xy) and orthogonal z-stack view (xz, yz) of the indicated area (white square) showing mitoGFP+ punta (white arrows) inside the HN12 mCherry-Ftractin receptor tumor cell (actin filaments, in magenta). Z-stacks were acquired in a sequential acquisition exciting first at 900 nm to visualize the mitoGFP (mitochondria in green) and second at 1080 nm to observe the mCherry-Ftractin (actin filaments here showed in magenta). Total thickness of 10.5 µm from 15 Z-stack in an interval 0.7 µm. Magnification 30x. Scale bar 20 µm.

Controlling cell density is essential for visualizing and quantifying TNTs. If cells are too far apart, TNT formation is impaired; if they are too close, the TNTs are too short to observe. Most current studies on TNTs use 2D substrates *in vitro*, which allow for easy control of cell density by adjusting cell concentrations or employing micropatterning to modify cell distances^41^. *In vivo*, this process is more complex. In our orthotopic model, HN12 cells form a tumor niche resembling human patient HNSCC tissue (Supplementary Fig. S6, Video S12 and S13). The high cell density in this niche physically restricts cell movement, limits migration, and hinders visualization of potential TNTs between cells (Supplementary Video S14).

Nevertheless, we successfully observed TNT-like connections *in vivo.* These connections exhibited structural features similar to those observed *in vitr*o, including composition (actin filaments), thickness (805 ± 108 nm), length (25-27 µm), and lifetime (∼2 hours) (Fig. 6B, Supplementary Video S15), as well as the presence of mitochondria inside the TNTs (Fig. 6C). Additionally, we visualized GFP+ puncta in Acc cells expressing F-tractin mCherry (Fig. 6D). While we were unable to monitor the complete transfer of mitochondria from D to Acc populations, the presence of mitochondria (mito-GFP+) in Acc cells and in TNT-like structures suggested that mitochondrial transfer may occur through these structures.

Furthermore, live imaging of the tumor in the mouse tongue revealed examples or both mechanisms responsible for TNT formation, similar to those described *in vitro*^22^: i)actin mediated protrusion and ii) cells dislodgement (Fig. 7A-B, Supplementary video S16 and S17, respectively). Since we visualized TNTs from two distinct cell populations HN12 mitoGFP-lifeActBFP2 (mitochondria green, actin blue) and HN12 F-tractin-mCherry (actin in red), excluding the possibility that these connections were derived from mitotic events between daughter cells (Supplementary Video S18). Taken together these data support the existence of TNT *in vivo* likely transferring mitochondria between tumor cells. Furthermore, due to the high cell density and the inherent challenges of imaging in living animals, we probably underestimated their prelevance.

**Fig. 7.**
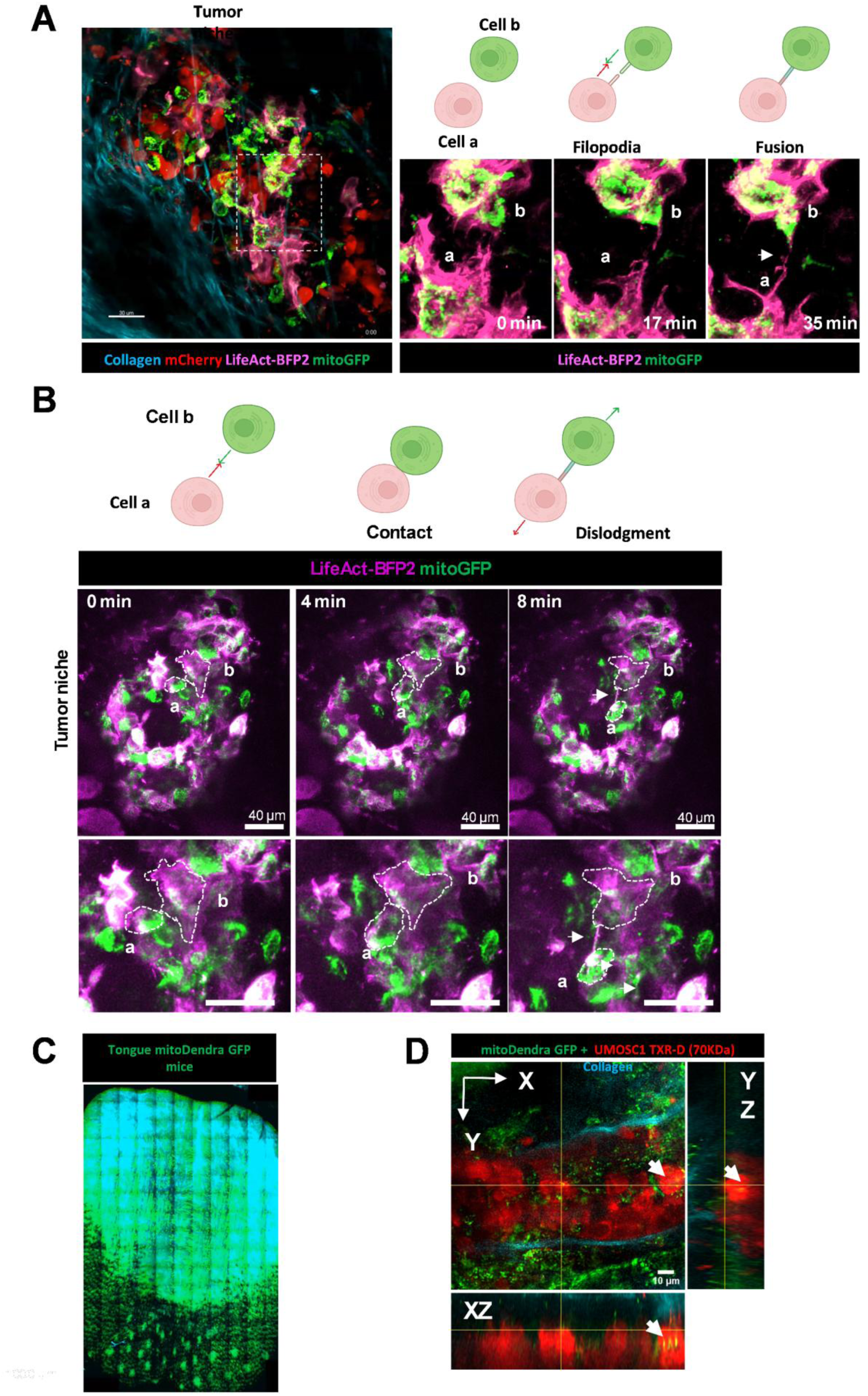
| TNT-like connection formation and mitochondria transfer to HNSCC from the TME *in vivo*. (A-B) Two different mechanisms of formation of TNT-like connections between HNSCC *in vivo*. (A) Representative time-lapse images from an intravital microscopy showing the formation of TNTs by Filopodia-growth mechanism inside the tumoral niche. Two different cells (a and b) grow filopodia that finally enter in contact. Note the presence of a mitoGFP dot inside the TNT, at the final time point (Supplementary video S16). (B) TNT formation by cell dislodgement mechanism: Representative images from an intravital microscopy movie at different time points with a magnification at the bottom for better visualization of the ROI showing two cells inside the tumoral niche that are in contact at initial time point and move in opposite directions generating a TNT between them (Supplementary video S17). Above each panel representative schema of the corresponding mechanism of formation. Dash areas indicate the ROI of interest. White arrows point to the presence of TNT. (C-D) HNSCC receiving mitochondria from the TME *in vivo*. Mito-Dendra1 mice were anesthetized, and the tongue was placed in the holder to minimize artifacts due to heartbeat and respiration. (C) Representative image of the whole tongue of the mito-Dendra1 mouse, showing how all host mitochondria (transgenic mouse) are green. The tongue image was obtained using 2-Photon microscopy, with an excitation wavelength of 500 nm and an emission filter of 505-560 nm to visualize GFP. Scanning was initiated from one of the anterior corners of the tongue using 3D stitching. Magnification 30x. Scale bar 1000µm. (D) Orthogonal reconstruction showing HNSCC mouse cell line 4MOSC1, that recapitulates human tobacco-related HNSCC genomic alteration and mutational landscape, labelled with TXR-70 (red) with host mitochondria inside, white arrows point the cell that contains GFP+ dots inside visible in both xy and xz views. Z-stacks were acquired exciting first at 750 nm to image collagen fibers (light blue), second at 900 nm to visualize the mitochondria (green) and finally at 1080 nm to acquire the TXR-70 4MOSC1 cells (tumor cells in red). Scale bar 10µm.

To assess whether healthy cells from the TME would transfer mitochondria to the tumor we used a syngeneic murine tongue model based on the HNSCC mouse cell line 4MOSC1^49^ orthotopically implanted into transgenic mice expressing the mitochondria-localized probe mito-Dendra-Tom^50^(Fig. 7C).4MOSC1 were labelled with Texas-RED (MW 70 kDa, TXR-D), which accumulates in the lysosomes and labels the cells for several days before injection into the tongue^51,52^. ISMic showed that TXR-70 HNSCC cells generated tumor in the tongue of mitoDendra1 mice, and we could observe tumor-Acc TXR + containing mitoDendra+ punta inside the cell body (Fig. 7D). This suggests that mitochondria from the TME cells were transferred to HNSCC tumor cells *in vivo*, serving as a proof-of-concept for monitoring this process in real time in live animals.

## Discussion

Communication between non-tumoral cells and tumors is a hallmark of cancer^53^. The interaction between tumor cells and the surrounding stromal components can drive tumor initiation^54^, progression^8,9^, and metastasis^55^. The horizontal transfer of mitochondria between different cell types in cancer or other diseases has been linked to tissue repair^56^, rescue mechanisms^12,15^, and enhanced tumorigenesis^4,5^ and metastasis^14,57^. Specifically, TNT-mediated transfer of healthy mitochondria has been associated with rescuing cells in early apoptotic stages^58^, correcting mitochondrial import failure^12^, and improving the metabolic fitness of T cells^7^. Moreover, various cancer types exploit TNTs to acquire mitochondria from immune cells^5^, mesenchymal stem cells (MSCs)^11^ and cancer-associated fibroblasts (CAFs)^9^ — supporting immune evasion^6^, promoting tumor cell survival, increasing malignancy, and driving invasion^59^. However, the occurrence of mitochondria transfer in cancer in the living animal, as well as the mechanisms and effects of mitochondrial exchange between the TME and cancer cells *in vitro* remain under characterized.

Here, we conducted a detailed live imaging analysis to study the physical characteristics of TNTs capable of carrying mitochondria, as well as the dynamics of TNT-mediated mitochondrial transfer between GB/GSCs, and between AS and GB/GSCs *in vitro.* Additionally, we used ISMic to assess TNT formation in a solid tumor model in live animals.

*In vitro* TNTs can be classified into two main types based on their cytoskeletal composition: (1) thin and fragile TNTs composed exclusively of actin filaments (Actin-based TNTs)^21^, and (2) thick TNTs composed of both actin filaments and MT (MT-based TNTs)^20,22^. Our findings reveal significant heterogeneity in the cytoskeletal composition of GB/GSCs, with thin Actin-based TNTs being more prevalent across all cell types. We demonstrate that mitochondria are transferred horizontally via both types of TNTs, providing evidence of mitochondria integration from the D population into the mitochondrial network of the recipient cell.

In neurons, short-range movement of mitochondria occurs via the actin cytoskeleton, whereas long-distance transport is MT-dependent^60^. Consistent with this, our findings show that MT-based TNT are longer and more stable compared to the Actin-based ones.

The central finding of our study is that mitochondrial transfer via TNTs serves as mode of cell-to-cell communication between GB/GSC tumor cells and the AS from the TME. We demonstrate the bidirectional interchange of mitochondria via TNTs, between primary non-tumoral AS and GB/GSCs. GB/GS cells deliver disrupted mitochondria to AS, which colocalize with astrocytic lysosomes suggesting a clearing mechanism through mitophagy. Conversely, AS shed healthy mitochondria to GB/GS cells, and these mitochondria likely integrate into the mitochondrial network of recipient cells. Our RNA bulk seq analysis of GSC containing AS-derived mitochondria reveals that transfer of heathy mitochondria (from AS to GSCs) through TNTs leads to the upregulation of genes related to mitochondrial metabolism (see model in Fig. 8). These findings suggest that the predominant functional impact results from the delivery of healthy mitochondria from AS to tumor cells, promoting increased expression of genes related to energy production and mitochondrial transport machinery.

**Fig. 8.**
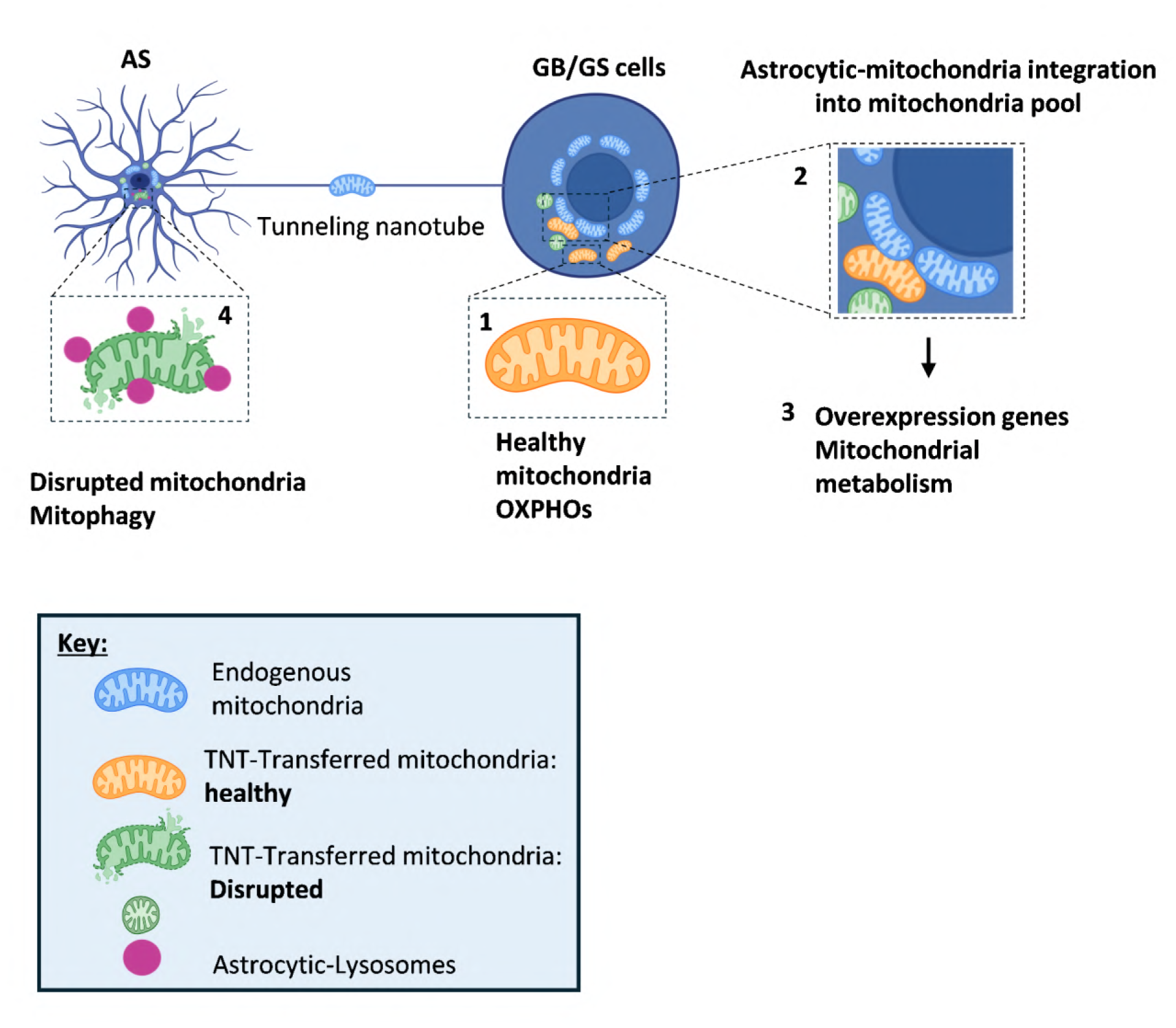
| Proposed model of mitochondrial interchange between GB/GS cells and AS via TNTs. **(1)** TNTs mediate the transfer of healthy mitochondria from AS to GB/GS cells and very few globular mitochondria, **(2)** these healthy mitochondria integrate into the GB/GS mitochondrial network, **(3)** leading to an increase in the expression of genes related to mitochondrial metabolism, and **(4)** TNTs also mediate the transfer of damaged mitochondria from GB/GS cells to AS, facilitating the clearance of damaged mitochondria by AS. Crated using BioRender.

Nonetheless, the reciprocal transfer of damaged mitochondria from GB/GS cells to AS may also contribute functionally—possibly by offloading dysfunctional organelles to evade apoptosis and improve tumor cell survival under stress conditions.

Several studies report that horizontal mitochondrial transfer enhances the metabolic capabilities of recipient cells^4,7,11,14–16,24^, while simultaneously depleting the D population, as observed in T cells^5^. Notably, the horizontal transfer of mutated mtDNA from tumor cells to T cells has been identified as a mechanism of immune evasion through complete endogenous mitochondrial replacement^6^.

Recently, Hoover et al. described the transfer of mitochondria from neurons to breast cancer cells, demonstrating that neuronal donation of mitochondria enhances tumor metabolic plasticity and promotes metastatic potential^14^. In addition, Neuzil’s group showed that the transfer of mitochondria from AS to mitochondrially impaired GBM cells restored mitochondrial respiration in GBM cells *in vivo*, enabling tumor initiation and growth in the brain^15^. These findings highlight the important role of TME-derived mitochondrial transfer in cancer progression.

Together, our data indicate that GSCs within the TME not only eliminate non-functional mitochondria and acquire healthy mitochondria by communicating with neighboring AS but are also stimulated to produce new active mitochondria, making them more metabolically active. These observations are consistent with previous reports in the central nervous system (CNS), where AS transfer healthy mitochondria to neurons following ischemic stroke as a neuroprotective effect^31^, or to GSCs, increasing GBM tumorigenicity^4,15^. Additionally, AS have been shown to degrade disrupted mitochondria from neurons in normal physiological conditions in the mouse optic nerve head *in vivo,* in a process called transmitophagy^61^. In Alzheimer’s disease, this neuron-AS transmitophagy increases, suggesting that TNTs may serve as a potential pathway for the transcellular transfer of disrupted mitochondria from neurons to AS^30^. Furthermore, studies outside the CNS have demonstrated that exogenous mitochondria derived from mesenchymal stromal cells (MSC) and acquired by endothelial cells trigger mitophagy rather than integrating into the endogenous mitochondrial pool^62^.

However, it is important to note that while we track the transfer of mitochondria, it is likely that other cellular materials, such as mRNA, microRNA, or lncRNAs^63^ are also transferred concomitantly.

The presence of TNTs in cancer has primarily been described *in vitro* or *ex-vivo* due to the difficulty of visualizing these thin and fragile connections with enough resolution in living animals. A breakthrough study from the Winkler’s group in 2015, showed the presence of long and thick intercellular connections, termed TM capable of transferring calcium waves, using two-Photon microscopy in live animals^27^. Recently, Watson and colleagues claimed the transfer of mitochondria from AS to GBM *in vivo* via TM. However these connections were identified *ex-vivo* after fixation^4^, and the distinction between TNTs and TMs remains unclear. We have previously identified both TMs and TNTs in tumor organoids derived from GSCs. Our live imaging data in this 3D *in vitro* model suggested that only thin TNT-like connections, and not TMs, are capable of transferring mitochondria^18^. Using a similar approach in the developing zebrafish embryo, we have recently shown that TNTs capable of transferring mitochondria exist in living organism^64^.

Here we used ISMic in an oral cancer model, which allowed for less invasive tumor observation compared to the GBM model, eliminating the need for a glass window on the skull^65^. This approach reduced infection risk and minimized imaging artifacts from animal respiration, while enabling the use of higher magnification objectives for enhanced image quality. Through this method, we identified TNT-like connections between HNSCC cells in live animals. Notably, these connections share key characteristics with TNTs observed in 2D *in vitro* cultures, including length, thickness, and persistence. Furthermore, we observed that TNTs between HNSCC cells *in vivo* can form through the same mechanisms identified *in vitro*: cell dislodgment and filopodia-mediated growth^22^. Additionally, the presence of mitochondria within their lumen supports their role in transfer. Our data indicate that tumor cells can acquire mitochondria both from other tumor cells and from the tumor microenvironment (TME), suggesting that mitochondrial exchange operates through similar mechanisms *in vivo* as observed *in vitro* with GBM cells, highlighting a critical mechanism of intercellular communication in the TME. This ISMic-based approach represents an initial step toward understanding TME-to-tumor TNT-mediated mitochondrial transfer *in vivo*. As a next step, a more clinically relevant model—such as the carcinogen-induced tongue cancer model—could be employed^49^.

Collectively, our findings provide compelling evidence of TNT-mediated mitochondria transfer between tumor cells and the TME both *in vitro* and *in vivo*, elucidating some of the functional consequences for recipient cells. These insights have important implications for cancer biology and lay the foundation for future studies and potential therapeutic strategies.

## Methods

### Cell culture

The human U251 GBM-derived cell line (09063001, Merk) was cultured in DMEM-F12 (1:1) medium (Gibco, 31330-038) supplemented with 10% fetal calf serum (FCS, Eurobio) and 1% penicillin/streptomycin (final concentration 100 U/mL; Thermo Fisher Scientific). GSCs were cultured in suspension in DMEM-F12 (Sigma) supplemented with B27 (50×, Gibco), N2 (100×, Gibco), and 20 ng/mL FGF-2 and EGF (Peprotech), as previously described^18^. HN12 HNSCC-derived cell line was cultured in DMEM supplemented with 10% FBS and 1% penicillin/streptomycin. 4MOSC1 syngeneic-derived cell line previously described^49^, was cultured in Keratinocyte medium supplemented with SFM Growth supplement, EGF (5ng/ml), choler toxin (5x10^11M) and 1% penicillin/streptomycin. All the cells were maintained at 37°C in 5% CO2 humidified incubators. Fresh medium was added, or passages were performed to the cell culture every 2–3 days. Absence of mycoplasma contamination was verified with MycoAlertTM Mycoplasma Detection Kit (Lonza). All methods were carried out in accordance with the approved guidelines of our institution.

### Murine primary AS preparation

Primary cortical AS were prepared from C57BL/6 wild-type or ROSAmT/mGmT mice from our in-house colony (Institut Pasteur, Paris, France) and used at postnatal days 0 to 3 (P0–P3), as previously described^66^. Briefly, following decapitation of the pups, the meninges were removed from the brain, and cortices were pooled before homogenization. Dissociated cortical cells (neocortex) were cultured in 75 cm² flasks coated with 5 µg/mL poly-D-lysine (PDL) in complete AS medium, consisting of DMEM-F12 (1:1) (Gibco) supplemented with 10% FCS (Eurobio) and 1% penicillin/streptomycin (Thermo Fisher). Cultures were maintained at 37°C in a humidified atmosphere containing 5% CO₂ until they reached confluency (7–10 days *in vitro*). Culture medium was refreshed every three days. To inhibit proliferation of other glial cells, a mixture of 5-fluoro-2′-deoxyuridine and uridine (10 µg/mL; Sigma-Aldrich) was added for one week. For experiments, confluent cultures were detached with trypsin and seeded onto PDL-coated multi-well plates. Before initiating experiments, AS cultures were characterized by immunostaining with rabbit anti-GFAP (1:500; Z0334 Dako), a well-established AS marker (see schematic, Figure S4 A).

### Lentiviral constructs

To label mitochondria, the pLV-CMV-mito-GFP (632432, Takara) and pLV-CMV-DsRED-mito (632421, Takara) plasmids were utilized. These plasmids encode subunit VIII of human cytochrome c oxidase fused with GFP or dsRED fluorescent proteins, respectively.

For labeling actin filaments in different colors, the following constructs were used: pLV-Ftractin-mCherry (#85131, Addgene), pLV-lifeact-mTagBFP2 (#101893, Addgene), and pLV-lifeact-iRFP670 (#84385, Addgene). Each construct was selected to provide distinct fluorescent markers to track actin dynamics in multicolor imaging applications.

For MT visualization, pLV-L304-EGFP-Tubulin-WT (#64060, Addgene) was employed. This plasmid encodes EGFP-tagged tubulin, allowing for the observation of MT structures within the cellular environment.

### Lentiviral particle production and transduction

Lentiviral particles (LVs) were generated in HEK 293T cells cultured in DMEM-F12 (Gibco), supplemented with 10% FCS (EuroBio) and 1% penicillin/streptomycin (Thermo Fisher) at 37°C in a humidified atmosphere containing 5% CO₂. The cells were seeded in T75 flasks the day prior to transfection to achieve 50–70% confluency. Transfection was performed using a mixture of plasmids encoding lentiviral components, specifically pCMVR8.74 (Gag-Pol-HIV1) and pMDG2 (VSV-G), along with the plasmid of interest at a ratio of 4:1:4 (μg), respectively. FuGENE HD Transfection reagent (Promega) was used according to the manufacturer’s protocol. After 48 hours, LVs were concentrated using LentiX-Concentrator (Takara Bio), and the resulting pellet was resuspended in 1 mL of PBS. For transduction, 500 µL of the resuspended LVs were added directly to the target cells of interest, including U251, GSLCs, HN12, or primary cortical AS, which had been plated in T75 flasks the day before infection to achieve 50–70% confluency. The cells were allowed to incubate with the LVs for a minimum of 48 hours for effective transduction. Positive cells were subsequently sorted using a BD FACS Aria III cell sorter (BD Biosciences).

### Live imaging

Two microscopes were used to perform time-lapse movies and analysis on live samples in 2D-classical culture: Spinning disk X1 Metamorph RCH (SD M) and Spinning disk W1 Nikon eclipse Ti2 confocal system (SD Ti2) (Nikon Instruments, Melville, NY, USA). These microscopes are equipped with a climate box to maintain humidity at 95%, temperature at 37°C, and 5% CO2 concentration. The utilized excitation laser was a white light laser (WLL), adjustable across wavelengths from 470 nm to 670 nm. To minimize cell stress, cell death, and photobleaching, the power of the WLL was maintained at the lowest possible level during imaging acquisition. All samples were acquired as z-stacks, with an average overall size of 7 µm, covering the whole volume of cells, either when they were alone or in co-culture, and a z-step size varying between 300 nm and 700 nm. The magnification used for all the mitochondria tracking movies and analysis of mitochondria health ranged from a minimum 40X to 100X, depending on the aim of the experiment.

Objective used:

– 60X oil immersion objective (Nikon APO60x NA=1.4 CSU) on SD M.
– 40X water immersion objective (Apo LWD 40X NA=1.15 WD=610 µm); and oil immersion objective 60x (Plan Apo 60XOIL NA=1.42 WD=150µm) and 100X (Plan Apo 100X OIL NA=1.45 WD=130 µm) on SD Ti2.

### Live-imaging analysis

#### a) Tracking mitochondrion movement inside the TNTs

The TrackMate program, developed by Jean-Yves Tinevez, was utilized to generate kymographs and analyze mitochondrial length and velocity within the TNTs over time, as detailed in the Data and Methods availability section of this publication in FIJI^67^. This tool can track objects inside a TNT while the connected cells migrate, providing the mitochondrial position relative to the TNT throughout the transfer process and yielding data on velocity and movement direction (anterograde, static, and retrograde).

Since the objects travel along a line following the TNT, assessing their speed is best accomplished via kymograph analysis. However, as the cells connected by the TNT move, the TNT itself changes position and size during imaging, making traditional kymograph tools inadequate. To address this, we developed a custom tool that constructs kymographs across a moving line over time. This tool relies on TrackMate^68^ to manually track the two endpoints of the TNT, and a custom extension enhances it by offering kymograph analysis and object tracking in the reference frame of the TNT.

The analyzed videos ranged from 20 minutes to 3 hours, depending on the cytoskeletal composition, with a 2-minute interval between frames. Minimal laser intensity was employed for each channel to avoid phototoxicity.

#### b) TNT Thickness/diameter over time

Most analyses of TNT thickness are conducted on fixed samples. However, TNTs are highly dynamic structures that are very sensitive to fixation. Furthermore, the fixation process can disrupt some of these connections, particularly those involving Actin-based TNTs.

In this study, TNT thickness was measured using live imaging assays with a custom Jython macro^67^ (see Data and Methods). Briefly, we assessed the intensity profile in the LifeAct fluorescent channel, fitting this profile with a Gaussian function to derive the TNT thickness. This approach allowed us to obtain TNT thickness measurements across different time frames of the recorded movie.

#### c) Mean-square displacement (MSD) of mitochondria

MSD was performed on the mitochondria tracked within the TNT, using a methodology and a MATLAB tool previously developed^38^. The dataset for the analysis included the tracks of 24 mitochondria. We fitted the first 25% of the log-log of the 24 MSD curves by a straight line and kept only the fits with a R^2^ larger than 0.8. The distribution of the slopes of the fits were larger than 1 (one-sided t-test, p < 10^-10^, confidence interval [1.58 - ∞]), revealing that the mitochondria are undergoing active motility inside TNTs.

#### d) Manders’ co-localization coefficients assay

The co-localization of D-derived mitoGFP with tetramethylrhodamine methyl ester (TMRM, 50 nM, active mitochondria) or Lysotracker Blue DND-22 (50 nM) in recipient cells was quantified using the JACoP plugin in FIJI, providing a measure of the overlap between two fluorescence signals, with values ranging from 0 (no overlap) to 1 (perfect overlap). After co-culturing D mitoGFP (mitochondria in green) and Acc lifeAct-670 cells (actin filaments in far-red) for 4 days, the double-positive cells were sorted, seeded overnight, and labeled with TMRM ± Lysotracker Blue. Images were acquired using SD at 37°C and 5% CO₂. For each individual cell, fluorescence channels were extracted, and thresholds were adjusted to calculate the Manders’ co-localization coefficients.

### Fixed samples imaging

Fixed samples were imaged using a laser scanning confocal microscope LSM 700 (Zeiss). Images were acquired with either a 40X or a 63X oil immersion objective (zoom 0.5), both of which have a numerical aperture of 1.4, using the Zen acquisition software (Zeiss). For each sample, a 3x3 mosaic image was captured with a 10% overlap to expand the field of view. The acquired images were then processed using ICY software (Quantitative Image Analysis Unit, Institut Pasteur, http://icy.bioimageanalysis.org) or FIJI software.

### TNT counting

TNTs were identified according to the protocol of Sáenz-de-Santa-María I. et al.^69^ Cells were plated at 50% confluence for TNT visualization, at a density of 40,000 to 60,000 cells/cm². The adhesion surface was pre-coated with either poly-D-lysine (0.1 mg/ml), for AS alone or AS-U251 co-culture, or laminin (10 mg/ml, Sigma) for AS-GSCs co-cultures. The percentage of TNT-connected cells was analyzed in live, unfixed samples to assess mitochondrial presence within TNTs and to evaluate cytoskeletal composition. For general quantification of TNT-connected cells, fixed samples were stained with the membrane marker Wheat Germ Agglutinin (WGA), which enhances visualization of these structures. To preserve TNT integrity during fixation, cells were treated with solution 1 (2% paraformaldehyde PFA, 0.05% glutaraldehyde, and 0.2 M HEPES in PBS) for 15 minutes, followed by an additional 15 minutes with solution 2 (4% PFA and 0.2 M HEPES in PBS) at 37°C.

### Quantification mitochondria transfer by Flow Cytometry Assays (FACS)

Transfer assays were performed according to the protocol of Sáenz-de-Santa-María I et al.^69^ Experimental triplicates were performed for each co-culture condition. To monitor transfer by secretion in a 2D co-culture setup, D and Acc cells were co-cultured with a 1 µm filter separating them. This filter prevents direct cell-cell contact while allowing the passage of mitochondria (0.5–1 µm in diameter). For FACS analysis, cells were passed through a cell strainer to remove aggregates and then fixed in 2% PFA. FACS data were acquired on a BD LSR Fortessa flow cytometer, with GFP, mCherry, and far-red fluorescence detected at 488 nm, 561 nm, and 670 nm excitation wavelengths, respectively. Ten thousand events were acquired per condition, and data were analyzed using FlowJo software.

### RNA-seq

GSC (+) have received astrocytic mitochondria (mito-dsRED+) and without astrocytic mitochondria (mito-dsRED-) were sorted after 4 days of co-culture in 2D-culture with murine AS expressing mito-dsRED in a ratio of 3 D to 1 Acc (GSC +), to allow the accumulation of mitochondria, from three independent experiments. After sorting, the cells were examined by fluorescence microscopy, and RNA was extracted using the RNeasy Micro Kit (Qiagen).

cDNA libraries were prepared using Illumina Stranded mRNA library Preparation Kit (Illumina) following the manufacturer’s protocol from 250 ng of RNA. Pooled libraries were additionally purified of unbound adaptors and primer dimers on AMPure XP magnetic beads (Beckman-Coulter). Sequencing was performed on a NextSeq2000 sequencing system (Illumina) to generate 40-60 million reads per sample (150bp).

The RNA-seq analysis was performed with Sequana^70^ version 0.16.11. In particular, we used the RNA-seq pipeline (v0.19.1) available on https://github.com/sequana/sequana_rnaseq, based on Snakemake framework^71^. The RNA-seq pipeline trimmed the reads from adapters and low quality bases using fastp software v0.20.1^72^. Then it mapped the reads to the Homo Sapiens genome (GRCh38 *hg38*) using STAR^73^. Note that we decided to perform a dual mapping by also including the Musculus genome because small contamination with murine AS samples were also present in the reads. For the read count and subsequent statistical analysis, mouse content was ignored. The read count matrix on Human annotation file was then used to identify differentially regulated genes. Differential expression testing was conducted using DESeq2 library 1.30.0^74^, scripts (also available in Sequana v0.16.11).

Finally, HTML reporting was available as the output of the Sequana RNA-seq pipeline. Parameters of the statistical analysis included the significance (Benjamini-Hochberg adjusted p-values, false discovery rate FDR < 0.05) and the effect size (fold-change) for each comparison considered. All software used within the RNAseq pipeline were containerized and are available within the Damona project (damona.readthedocs.io) and downloaded automatically by the RNA-seq pipeline (enforcing reproducibility of the analysis).

Concerning the comparison of GSC (+) pos (mito-dsRED+) versus GSC (+) neg (mito-dsRED-), genes with an adjusted P value of <0.05 and log2 fold change >1 were selected for an enrichment analysis. We performed GO term enrichment using the Panther database^75^. These enrichments were performed with the Sequana standalone called enrichment-panther feeding the list of differentiated genes of interest. The Panther databases was accessed through the BioServices package^76^.

### Animal experimentation

All experiments were approved by the National Institute of Cancer Animal Care and Use Committee (NCI/NIH). The mice were housed in appropriate sterile filter-capped cages and provided food and water ad libitum.

#### a) Tongue tumor xenograft in immunocompromised athymyc nu/nu mice

The xenograft tumor generation was performed as previous described by Amornphimoltham et al^48^. Briefly, athymic nude mice were anesthetized using isoflurane 2–5% and maintained under continuous anesthesia through a nose cone. The tip of the tongue was gently pulled out of the mouth. Half a million HN12 cells, consisting of D cells co-expressing mitoGFP-lifeAct-BFP2 and Acc cells expressing F-tractin-mCherry in 20 µl saline, were submucosally injected into the lateral anterior region of the tongue using an insulin syringe. The needle was inserted up to 1–2 mm, injecting the cells as close to the surface as possible. The animals were fed a soft dough diet from the day of injection. Tumor growth was monitored, and ISMic was performed every two days for up to 40 days (Figure 6A). ISMic was conducted when the tumor reached 150 µm from the surface, allowing the acquisition of high-quality images.

#### b) Syngeneic HNSCC mice model

4MOSC1 derived-cell line, previous described to recapitulates the human tobacco-related HNSCC mutanome^49^, were labeled with dextran red (TXR-D, MW70 KDa) and injected (500, 000 cells/20µl saline) into the ventral tongue of mitoDendra2 mice, which express all mitochondria in green (B6;129S-*Gt(ROSA)26Sor^tm1.1(CAG-COX8A/Dendra2)Dcc^*/J, https://www.jax.org/strain/018397#, The Jackson Laboratory). After the development of the tumor ISMic were performed in the animal alive to visualize the transfer from the TME into de tumoral 4MOSC1 cells.

#### c) Intravital two-photon microscopy in rodents alive (ISMic)

After the tumor developed, we performed ISMic as previous described ^35^. Mice were anesthetized with intraperitoneal injections of ketamine (100 mg/kg) and xylazine (10 mg/kg). Animals were placed on a preheated stage with the mouth open, and the tongue was gently retracted using nontoothed forceps and held in a custom-made holder, secured with a glass coverslip. Body temperature was maintained at 37-38°C using a preheated stage. Imaging was performed by using an inverted laser-scanning two-photon microscope (MPE-RS, Olympus, Center Valley, PA, USA) equipped with a tunable laser (Insight DS+, Spectra Physics, Santa Clara, CA, USA). Excitation wavelengths of 750 nm, 900 nm, or 1080 nm were used to visualize collagen fibers, mitochondria, and F-tractin mCherry or TXR-70 Emitted light was collected by an appropriate set of mirrors and filters on 3 GaAsP detectors (bandpass filters: Blue = 410–460 nm, Green = 495–540 nm, Red = 575–645 nm). Images were acquired using 37°C heated objectives: 30X and 40X silicone oil immersion UPLSAPO objectives (NA 1.05 and 1.25 respectively), from Olympus. The first detector collagen fibers through SHG. The second captured mitoGFP and lifeAct-BFP2 signals, while the third detected mCherry and TXR-70. For time-lapse imaging of the live animals, the acquisition speed was set to 2 or 5 min of intervals, and duration up to 2 hours.

### Imaging post-acquisition procession: Deconvolution and 3D-rendering

Background noise in ISMic movies was reduced by applying a 2 x 2 pixel low-pass filter to each image for one or two rounds using Metamorph software (Molecular Devices). Images were then deconvolved using the Huygens Professional Deconvolution program (Scientific Volume Imaging B.V., Hilversum, Netherlands), which calculated a theoretical point spread function (PSF) based on the microscopy parameters. Deconvolution was performed with an interactive Classical Maximum Likelihood Estimation (CMLE) algorithm. For 3D volume rendering, Imaris 9.8.2 64-bit (Bitplane) was used. Drift correction was applied using the Correct 3D Drift plugin in FIJI. Final preparation of movies and images was managed with FIJI software.

### Statistical analysis

Statistical analyses and graphs were generated using the GraphPad Prism version 9 software. All the results are expressed as the mean ± SD. For comparisons between two groups the student’s t test was used. Unless stated in the figure’s legend, for comparisons between more than two groups, Two-way ANOVA with Tukey’s post hoc analysis was employed. In all cases, statistical significance was attributed when p ≤ 0.05.

## Data and methods availability

The data, tools, and material (or their source) that support the article are available from the corresponding author upon reasonable request. The tool to track mitochondria in TNTs is publicly available as a TrackMate extension, in the FIJI software platform. It can be freely installed by subscribing to the *TrackMate-Kymograph* update site in the FIJI updater.

## Acknowledgements

The authors thank Dr. Nicolas Melis, Dr. Thomas Madsen, and Dr. Yeap Ng at the National Institutes of Health (NIH) for their valuable discussions and invaluable assistance. We also thank R. Bouyssie from the administrative staff of the Membrane Traffic and Pathogenesis department at the Institut Pasteur (IP). Additionally, we would like to express our gratitude to Pierre-Henri Commere at Cytometry plataform IP in sorting cells, and to Laure Lemée for her quality control of the RNA-seq data.

## Author contributions

I.SSM. conceived the project, designed and executed the experiments, analyzed the data, prepared the figures, and wrote the manuscript. C.B. assisted with manuscript writing and discussion. J-Y. T. developed the TrackMate program and provided training for dynamic image analysis algorithms. E.MJC. provided the patient-derived GSCs. I.V. and T.C. provided the RNA bulk sequencing and analysis, helping I.SSM. with the interpretation of the data. R.W. guided and assisted with the conceptualization, resources, and supervision of the *in vivo* studies and contributed to manuscript writing. C.Z. conceived and supervised the project, guided manuscript writing, and provided the resources. All authors read, edited, and approved of the article.

## Extended Data

**Supplementary video S1:** Mitochondria transfer via TNT from U251-to-U251 in 2D-culture in vitro. U251 expressing mito-dsRED-lifeAct-670 (mitochondria in red and actin filament cyan). The total duration of the video is 2 hours and 18 min.

**Supplementary video S2:** Representative video showing the procedure for measuring TNT thickness over time using live imaging. Magnified ROI shows TNT connections between U251 cells expressing mito-dsRED and LifeAct-670 (mitochondria in red, actin filaments in cyan). A dashed line highlights the region where thickness measurements are performed throughout the video, avoiding the gondola-like structures filled with mitochondria. Total duration: 32 minutes, with 2-minute intervals.

**Supplementary video S3:** TNT between U251 mitoGFP-LifeAct-BFP2 (mitochondria and actin filaments in green and cyan, respectively) and U251 LifeAct-670 (actin filaments in magenta). A rare case where the TNT connection persists for up to 7 hours. Images were captured at 2-minute intervals. Magnification on the right provides a clearer view of the TNT connection.

**Supplementary video S4:** TNT between U251 mitoGFP-LifeAct-BFP2 (D cell, mitochondria and actin filaments in green and cyan, respectively) and U251 LifeAct-670 (Acc cell, actin filaments in magenta). The TL Phase channel is shown on the right. Note the presence of a mitoGFP+ dot inside the Acc cell. Total duration: 1 hour and 10 minutes, with 5-minute intervals.

**Supplementary video S5:** U251 mito-dsRED-lifeAct-BFP2-GFP-tubulin (mitochondria in red, actin filaments in blue, and microtubules in green) undergoing mitosis in a 2D culture. The video highlights Actin-based TNTs connecting mitotic cells with their neighboring cells.

**Supplementary video S6:** Tracking mitochondria traveling via Actin-based TNT in U251 mito-dsRED-lifeAct-BFP2-GFP-tubulin (mitochondria, actin filaments and tubulin in red, blue and green). On the right magnification of the TNT for better visualization of the tracking and TNT composition. The TL Phase channel is used to visualize cell membranes. Total duration: 24 minutes, with 2-minute intervals.

**Supplementary video S7:** Complete transfer of mitochondria via Actin-based TNTs in GSC (-) cells, labeled with mitoTracker Red (mitochondria in red), Actin Cell Mask 670 (actin filaments in blue), and SPY55-tubulin (tubulin in green). Total duration: 54 minutes, with 2-minute intervals.

**Supplementary video S8**: Tracking mitochondria traveling via MT-based TNT in U251 mito-dsRED-lifeAct-BFP2-GFP-tubulin (mitochondria, actin filaments and tubulin in red, blue and green). On the right magnification of the TNT for better visualization of the tracking and TNT composition. Total duration is 1 hour and 2 minutes, intervals 2 minutes.

**Supplementary video S9:** Fusion of mitochondria traveling via Actin-based TNT into the recipient cell’s mitochondrial network. Total duration: 58 minutes, with 2-minute intervals.

**Supplementary video S10:** Fusion of mitochondria traveling via MT-based TNT into the recipient cell’s mitochondrial network. Total duration: 1 hour and 14 minutes, with 2-minute intervals.

**Supplementary video S11:** 3D-rendering using Imaris of HN12 Ftractin-mCherry TNT in 2D-culture *in vitro*. Showing the TNT hovering in the medium.

**Supplementary video S12:** Z-stack image of the tumoral niche in an HNSCC xenograft model, showing HN12 mitoGFP-lifeActBFP2 (D cells) and HN12 F-tractin mCherry (Acc cells) in direct contact, acquired by ISMic on the tongue of a live rodent.

**Supplementary video S13:** 3D rendering of the previous tumor niche for better visualization of cell density and compaction. The tumoral niche closely resembles those found in HNSCC tissue samples.

**Supplementary video S14:** ISMic video of a tumoral niche formed by co-culture of HN12 labeled with TXR-70 and HN12 labeled with Cell Tracker Green, showing the inability of HN12 cells to migrate.

**Supplementary video S15:** Presence of TNT *in vivo*. Representative ISMic video showing the presence of TNT between HN12 mitoGFP-lifeAct-BFP2 cells in the tongue of a live animal. On the left, a global image displays the presence of D cells (mitoGFP-lifeAct-BFP2, with mitochondria and actin filaments in green and blue, respectively) and Acc cells (F-tractin mCherry, red) within the tumoral niche. Total duration: 1 hour and 50 minutes, with 5-minute intervals.

**Supplementary video S16:** TNT formation via filopodia growth mechanism *in vivo*. HN12 mitoGFP-lifeAct-BFP2 (mitochondria and actin filaments in green and magenta, respectively) forms TNT-like connections within a space in the tumor niche. On the left, the video shows all channels, allowing visualization of the tumor niche and collagen fibers (light blue). On the right, the video displays only the mitoGFP-lifeAct channels for enhanced visualization of TNT formation.

**Supplementary Video S17:** TNT formation via cell dislodgement mechanism *in vivo*. The tumor niche is formed by HN12 mitoGFP-lifeAct-BFP2 (mitochondria and actin filaments in green and cyan, respectively) and HN12 mCherry-F-tractin (actin filaments in red), with a physical space in between. The video shows the contact between two different cells, followed by their subsequent migration in opposite directions, forming a TNT between HN12 cells expressing mitoGFP-lifeAct-BFP2 (green and blue). Total duration: 32 minutes, continuous intervals.

**Supplementary Video S18**: *In vivo* heterotypic TNT connections between HN12 lifeAct-BFP2 and HN12 F-tractin-mCherry. 3D rendering and movie created with Imaris, showing a TNT-like connection between HN12 mitoGFP-lifeAct-BFP2 and HN12 F-tractin mCherry.

---

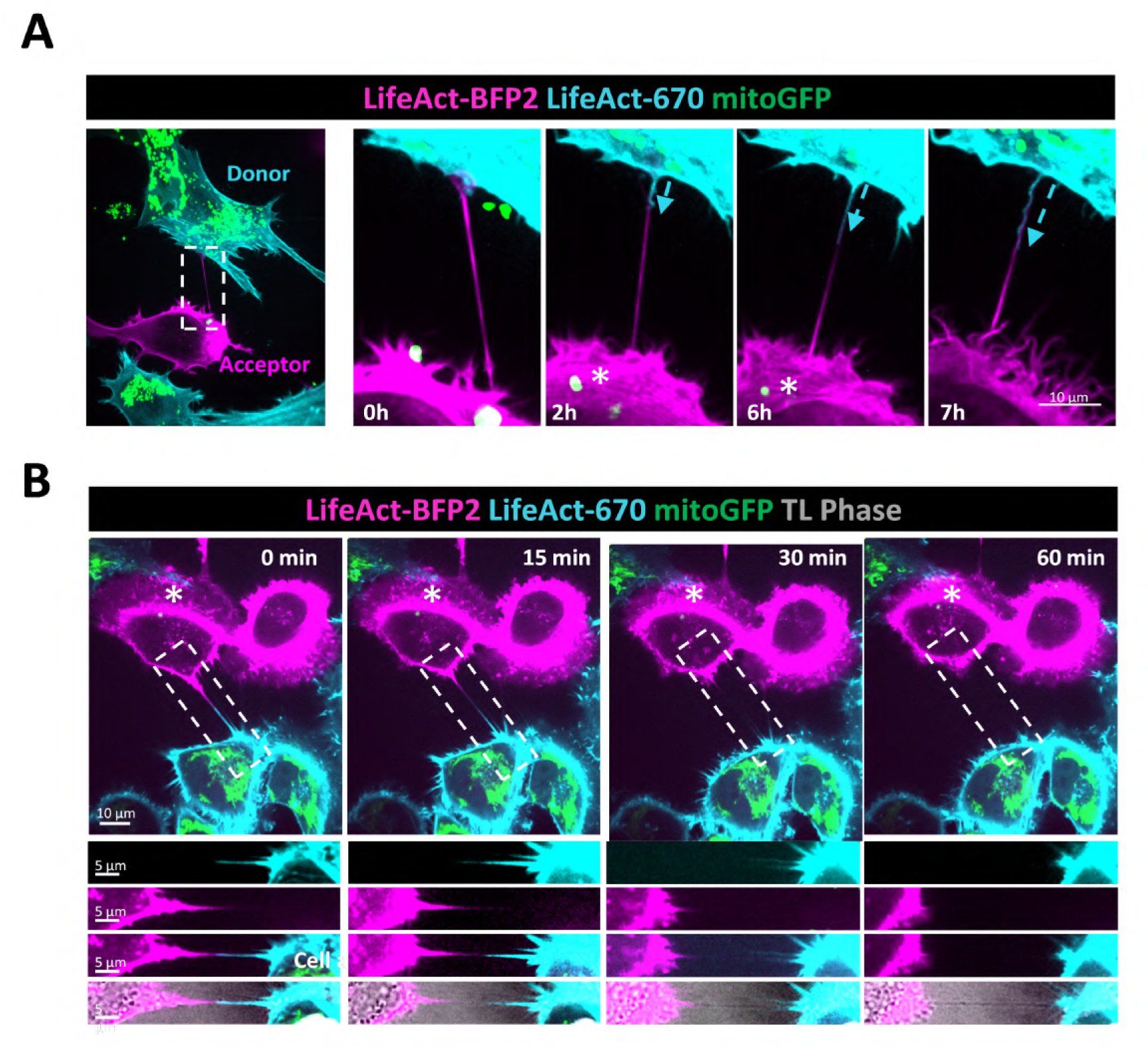

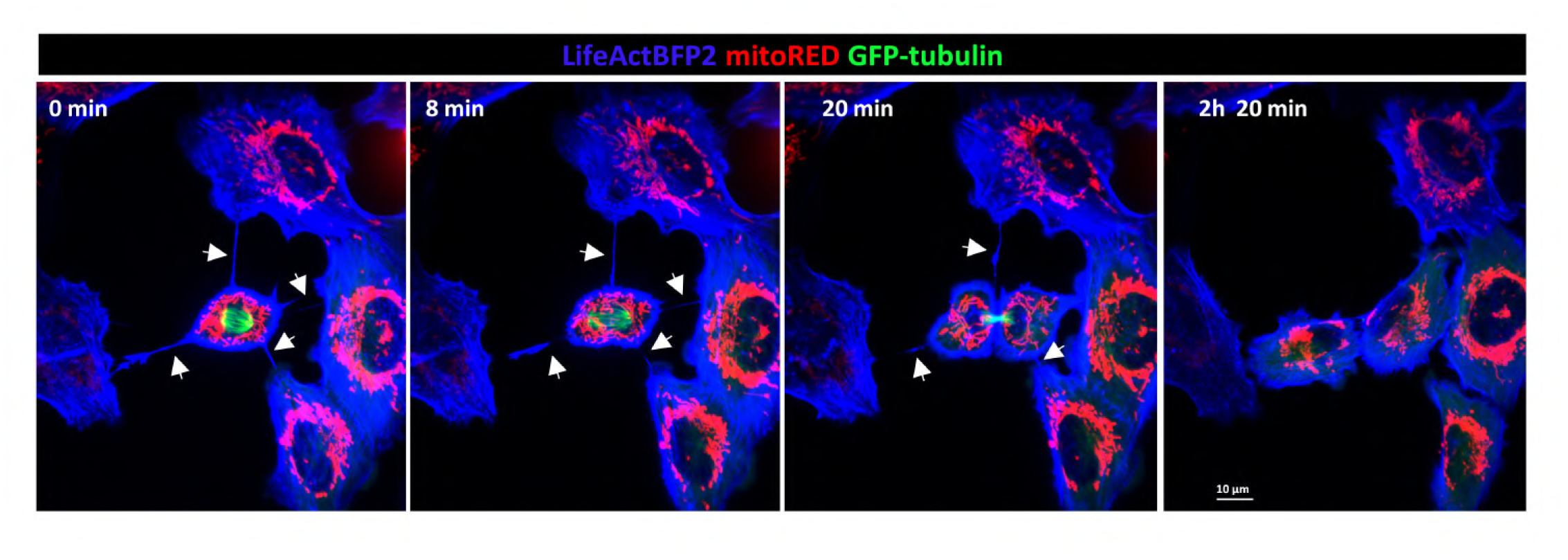

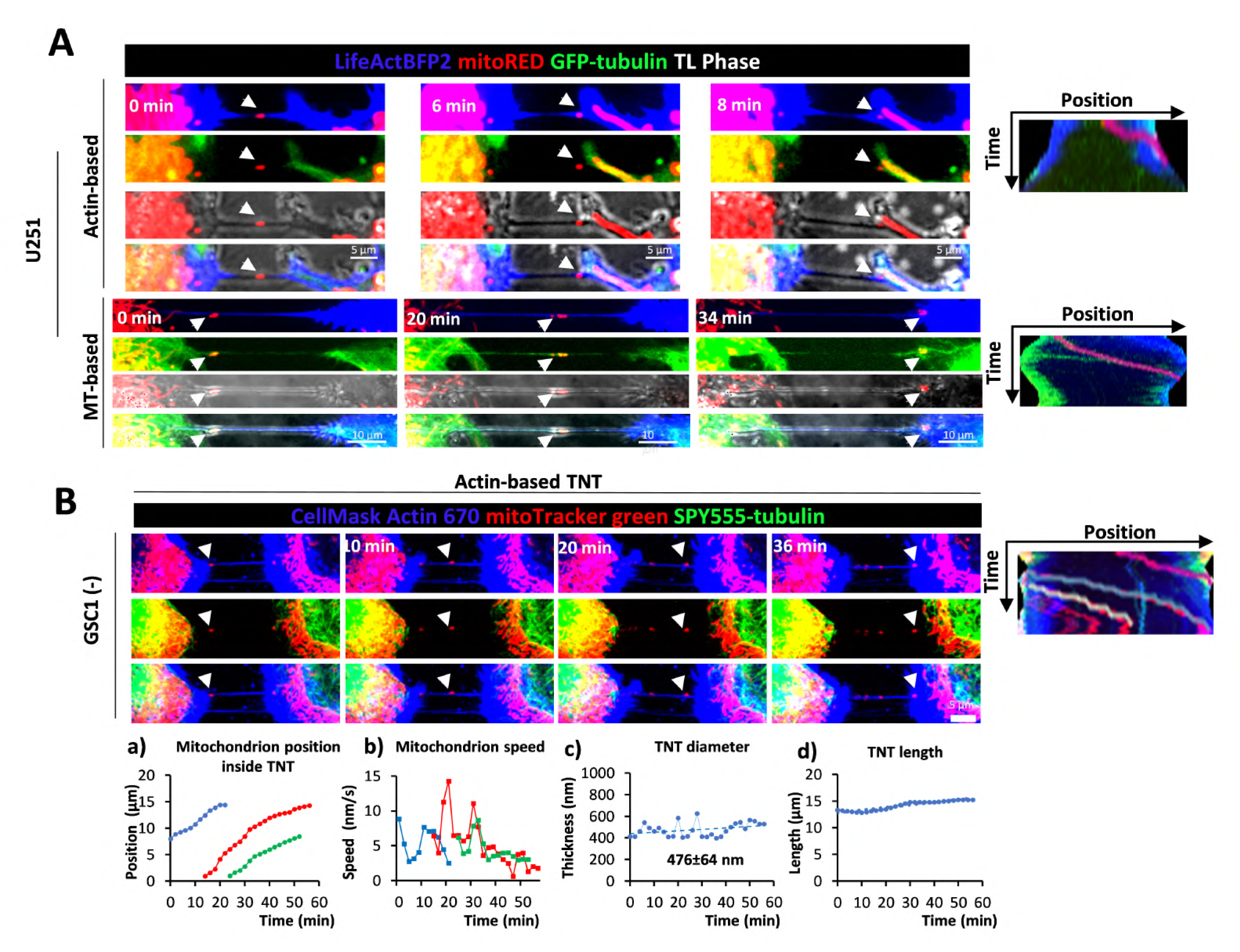

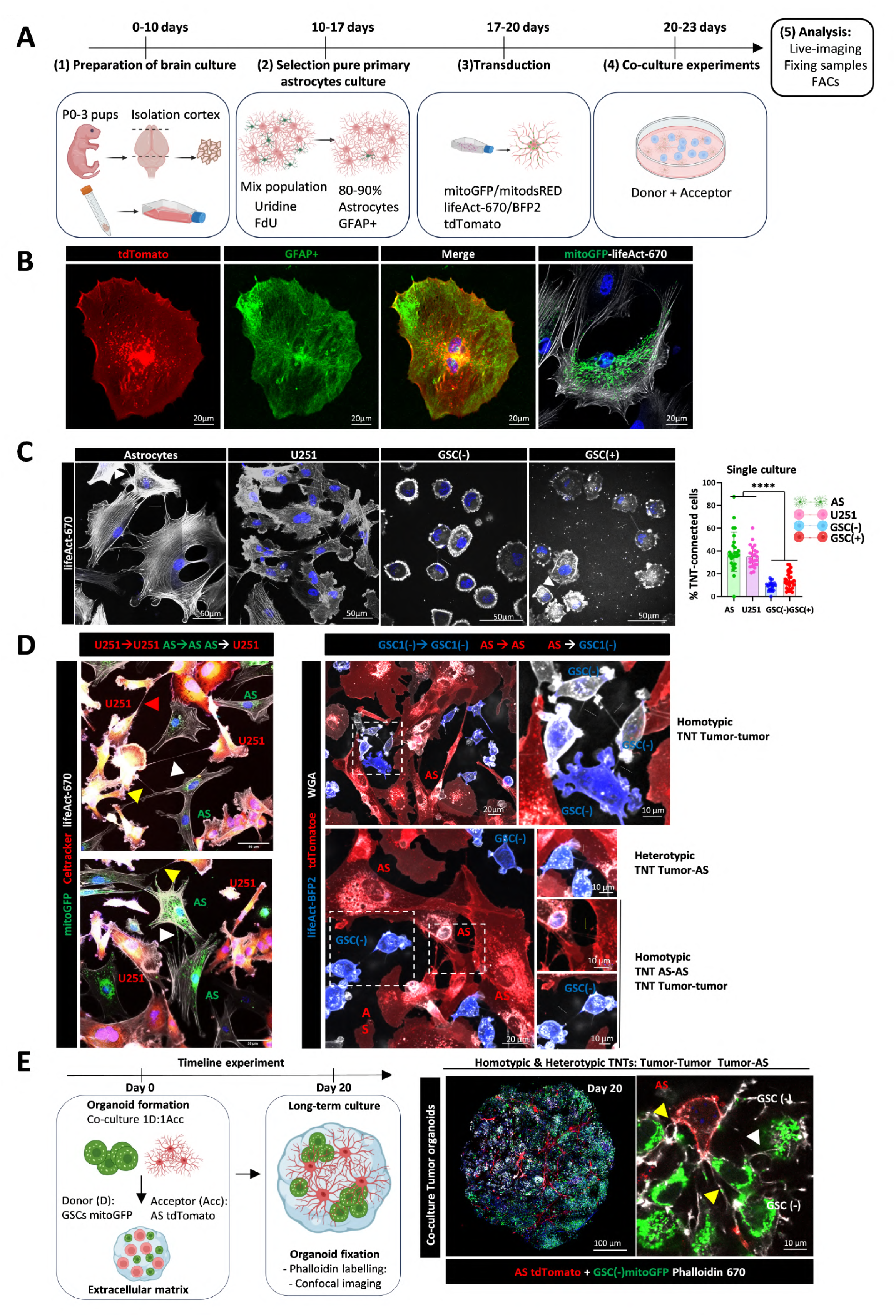

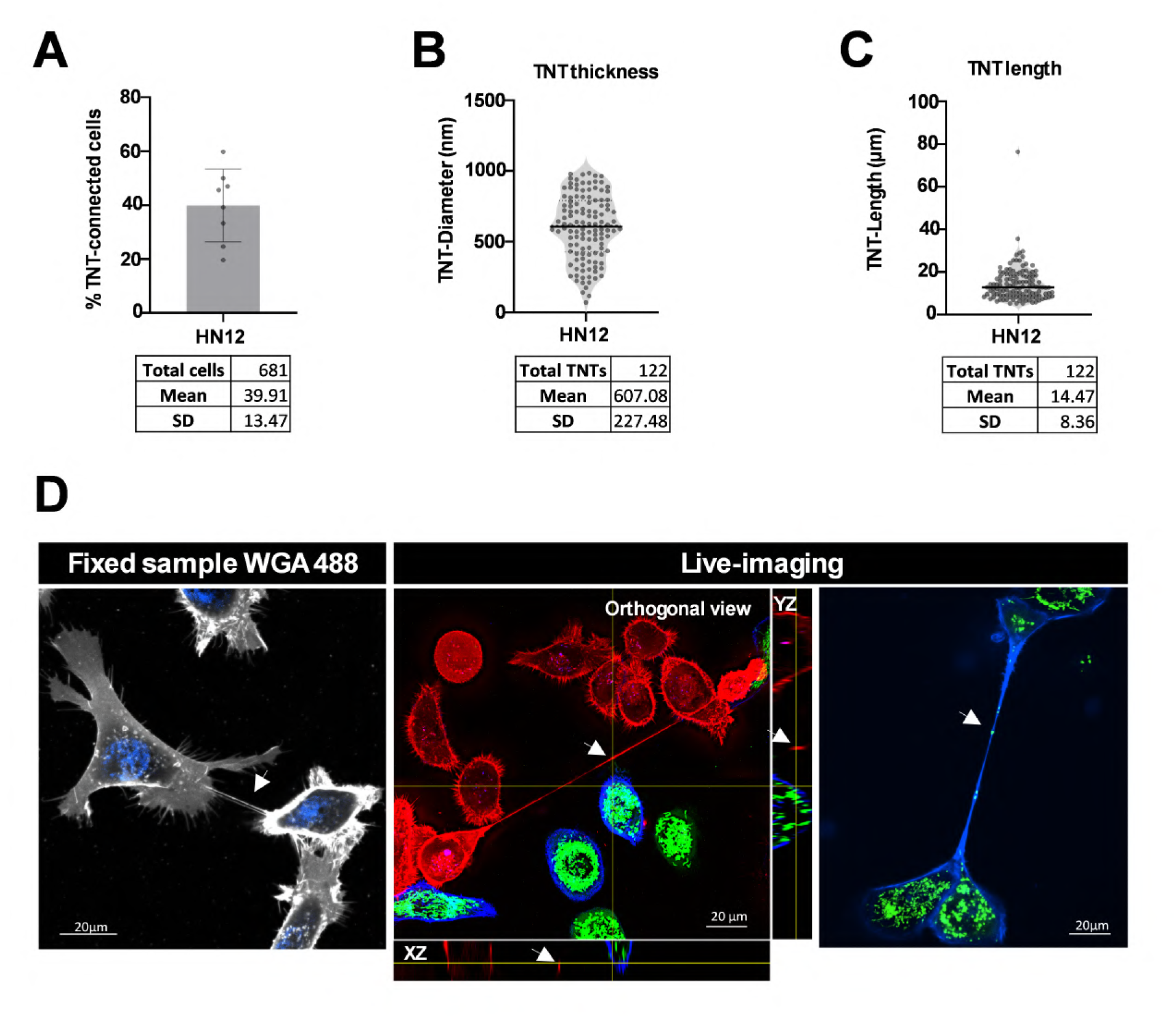

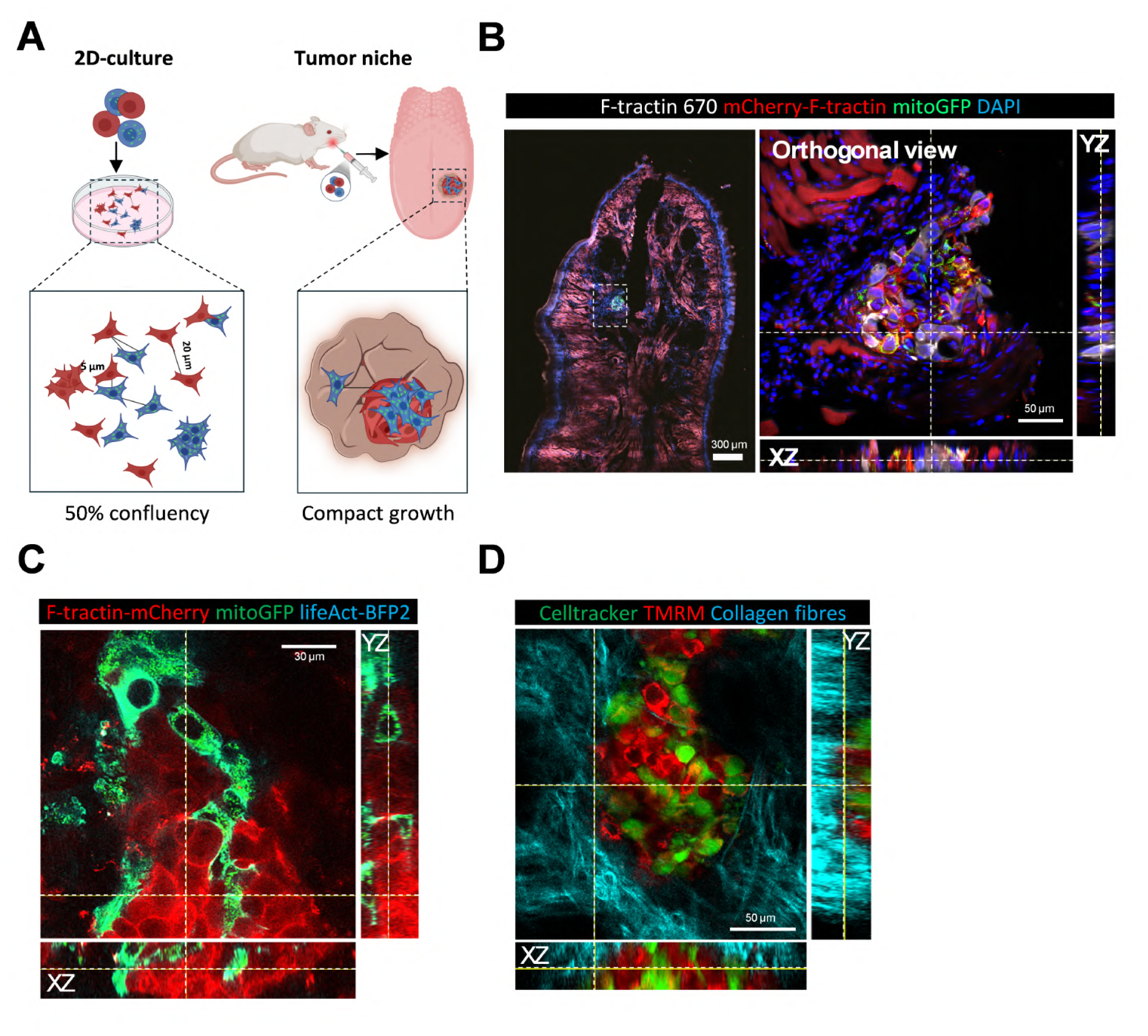

## References

1. Fontana, F. & Limonta, P. The multifaceted roles of mitochondria at the crossroads of cell life and death in cancer. Free Radical Biology and Medicine 176, 203–221 (2021).

2. Palma, F. R. et al. ROS production by mitochondria: function or dysfunction? Oncogene 43, 295–303 (2024).

3. Iorio, R., Petricca, S., Mattei, V. & Delle Monache, S. Horizontal mitochondrial transfer as a novel bioenergetic tool for mesenchymal stromal/stem cells: molecular mechanisms and therapeutic potential in a variety of diseases. Journal of Translational Medicine 22, 491 (2024).

4. Watson, D. C. et al. GAP43-dependent mitochondria transfer from astrocytes enhances glioblastoma tumorigenicity. Nat Cancer 1–17 (2023) doi:10.1038/s43018-023-00556-5.

5. Saha, T. et al. Intercellular nanotubes mediate mitochondrial trafficking between cancer and immune cells. Nat. Nanotechnol. 1–9 (2021) doi:10.1038/s41565-021-01000-4.

6. Ikeda, H. et al. Immune evasion through mitochondrial transfer in the tumour microenvironment. Nature 638, 225–236 (2025).

7. Baldwin, J. G. et al. Intercellular nanotube-mediated mitochondrial transfer enhances T cell metabolic fitness and antitumor efficacy. Cell 187, 6614–6630.e21 (2024).

8. Ippolito, L. et al. Cancer-associated fibroblasts promote prostate cancer malignancy via metabolic rewiring and mitochondrial transfer. Oncogene 38, 5339–5355 (2019).

9. Goliwas, K. F. et al. Mitochondrial transfer from cancer-associated fibroblasts increases migration in aggressive breast cancer. J Cell Sci 136, jcs260419 (2023).

10. Hekmatshoar, Y., Nakhle, J., Galloni, M. & Vignais, M.-L. The role of metabolism and tunneling nanotube-mediated intercellular mitochondria exchange in cancer drug resistance. Biochemical Journal 475, 2305–2328 (2018).

11. Nakhle, J. et al. Mitochondria Transfer from Mesenchymal Stem Cells Confers Chemoresistance to Glioblastoma Stem Cells through Metabolic Rewiring. Cancer Res Commun 3, 1041–1056 (2023).

12. Needs, H. I. et al. Rescue of mitochondrial import failure by intercellular organellar transfer. Nat Commun 15, 988 (2024).

13. Wang, X. & Gerdes, H.-H. Transfer of mitochondria via tunneling nanotubes rescues apoptotic PC12 cells. Cell Death Differ 22, 1181–1191 (2015).

14. Hoover, G. et al. Nerve-to-cancer transfer of mitochondria during cancer metastasis. Nature (2025) doi:10.1038/s41586-025-09176-8.

15. Brisudova, P. et al. Functional mitochondrial respiration is essential for glioblastoma tumour growth. Oncogene (2025) doi:10.1038/s41388-025-03429-6.

16. Berridge, M. V. et al. Horizontal mitochondrial transfer in cancer biology: Potential clinical relevance. Cancer Cell 43, 803–807 (2025).

17. Rustom, A., Saffrich, R., Markovic, I., Walther, P. & Gerdes, H.-H. Nanotubular Highways for Intercellular Organelle Transport. Science 303, 1007–1010 (2004).

18. Pinto, G. et al. Patient-derived glioblastoma stem cells transfer mitochondria through tunneling nanotubes in tumor organoids. Biochem J 478, 21–39 (2021).

19. Vignais, M.-L., Caicedo, A., Brondello, J.-M. & Jorgensen, C. Cell Connections by Tunneling Nanotubes: Effects of Mitochondrial Trafficking on Target Cell Metabolism, Homeostasis, and Response to Therapy. Stem Cells Int 2017, 6917941 (2017).

20. Sáenz-de-Santa-María, I. et al. Control of long-distance cell-to-cell communication and autophagosome transfer in squamous cell carcinoma via tunneling nanotubes. Oncotarget 8, 20939–20960 (2017).

21. Sartori-Rupp, A. et al. Correlative cryo-electron microscopy reveals the structure of TNTs in neuronal cells. Nat Commun 10, 342 (2019).

22. Cordero Cervantes, D. & Zurzolo, C. Peering into tunneling nanotubes-The path forward. EMBO J 40, e105789 (2021).

23. Babenko, V. A. et al. Miro1 Enhances Mitochondria Transfer from Multipotent Mesenchymal Stem Cells (MMSC) to Neural Cells and Improves the Efficacy of Cell Recovery. Molecules 23, 687 (2018).

24. Marlein, C. R. et al. CD38-Driven Mitochondrial Trafficking Promotes Bioenergetic Plasticity in Multiple Myeloma. Cancer Res 79, 2285–2297 (2019).

25. Valdebenito, S. et al. Tunneling nanotubes, TNT, communicate glioblastoma with surrounding non-tumor astrocytes to adapt them to hypoxic and metabolic tumor conditions. Sci Rep 11, 14556 (2021).

26. Yalamarty, S. S. K. et al. Mechanisms of Resistance and Current Treatment Options for Glioblastoma Multiforme (GBM). Cancers 15, 2116 (2023).

27. Osswald, M. et al. Brain tumour cells interconnect to a functional and resistant network. Nature 528, 93–98 (2015).

28. Bikfalvi, A. et al. Challenges in glioblastoma research: focus on the tumor microenvironment. Trends in Cancer 9, 9–27 (2023).

29. Read, R. D., Tapp, Z. M., Rajappa, P. & Hambardzumyan, D. Glioblastoma microenvironment-from biology to therapy. Genes Dev 38, 360–379 (2024).

30. Lampinen, R. et al. Neuron-astrocyte transmitophagy is altered in Alzheimer’s disease. Neurobiology of Disease 170, 105753 (2022).

31. Hayakawa, K. et al. Transfer of mitochondria from astrocytes to neurons after stroke. Nature 535, 551–555 (2016).

32. Lemarié, A. et al. The STEMRI trial: Magnetic resonance spectroscopy imaging can define tumor areas enriched in glioblastoma stem-like cells. Science Advances 9, eadi0114 (2023).

33. Ebrahim, S. & Weigert, R. Intravital microscopy in mammalian multicellular organisms. Curr Opin Cell Biol 59, 97–103 (2019).

34. Weigert, R., Sramkova, M., Parente, L., Amornphimoltham, P. & Masedunskas, A. Intravital microscopy: a novel tool to study cell biology in living animals. Histochem Cell Biol 133, 481–491 (2010).

35. Amornphimoltham, P. et al. Rab25 regulates invasion and metastasis in head and neck cancer. Clin Cancer Res 19, 1375–1388 (2013).

36. Pinto, G., Brou, C. & Zurzolo, C. Tunneling Nanotubes: The Fuel of Tumor Progression? Trends in Cancer 6, 874–888 (2020).

37. Matejka, N., Amarlou, A., Neubauer, J., Rudigkeit, S. & Reindl, J. High-Resolution Microscopic Characterization of Tunneling Nanotubes in Living U87 MG and LN229 Glioblastoma Cells. Cells 13, 464 (2024).

38. Tarantino, N. et al. TNF and IL-1 exhibit distinct ubiquitin requirements for inducing NEMO–IKK supramolecular structures. Journal of Cell Biology 204, 231–245 (2014).

39. Halász, H., Tárnai, V., Matkó, J., Nyitrai, M. & Szabó-Meleg, E. Cooperation of Various Cytoskeletal Components Orchestrates Intercellular Spread of Mitochondria between B-Lymphoma Cells through Tunnelling Nanotubes. Cells 13, 607 (2024).

40. Veranič, P. et al. Different Types of Cell-to-Cell Connections Mediated by Nanotubular Structures. Biophys J 95, 4416–4425 (2008).

41. Henderson, J. M. et al. Tunnelling nanotube formation is driven by Eps8/IRSp53-dependent linear actin polymerization. The EMBO Journal 42, e113761 (2023).

42. Notario Manzano, R., et al. Proteomic landscape of tunneling nanotubes reveals CD9 and CD81 tetraspanins as key regulators. Elife 13, RP99172 (2024).

43. Neikirk, K. et al. MitoTracker: A useful tool in need of better alternatives. European Journal of Cell Biology 102, 151371 (2023).

44. Desai, S. et al. Performance of TMRM and Mitotrackers in mitochondrial morphofunctional analysis of primary human skin fibroblasts. Biochimica et Biophysica Acta (BBA) - Bioenergetics 1865, 149027 (2024).

45. Kalargyrou, A. A. et al. Nanotube-like processes facilitate material transfer between photoreceptors. EMBO Rep 22, e53732 (2021).

46. Valdebenito, S., Audia, A., Bhat, K. P. L., Okafo, G. & Eugenin, E. A. Tunneling Nanotubes Mediate Adaptation of Glioblastoma Cells to Temozolomide and Ionizing Radiation Treatment. iScience 23, 101450 (2020).

47. Masedunskas, A., Porat-Shliom, N. & Weigert, R. Regulated Exocytosis: Novel Insights from Intravital Microscopy. Traffic 13, 627–634 (2012).

48. Amornphimoltham, P., Thompson, J., Melis, N. & Weigert, R. Non-Invasive Intravital Imaging of Head and Neck Squamous Cell Carcinomas in Live Mice. Methods 128, 3–11 (2017).

49. Wang, Z. et al. Syngeneic animal models of tobacco-associated oral cancer reveal the activity of in situ anti-CTLA-4. Nat Commun 10, 5546 (2019).

50. Porat-Shliom, N., Harding, O. J., Malec, L., Narayan, K. & Weigert, R. Mitochondrial Populations Exhibit Differential Dynamic Responses to Increased Energy Demand during Exocytosis In Vivo. iScience 11, 440–449 (2019).

51. Masedunskas, A. & Weigert, R. Intravital two-photon microscopy for studying the uptake and trafficking of fluorescently conjugated molecules in live rodents. Traffic 9, 1801–1810 (2008).

52. Madsen, D. H. et al. M2-like macrophages are responsible for collagen degradation through a mannose receptor-mediated pathway. J Cell Biol 202, 951–966 (2013).

53. Hanahan, D. & Weinberg, R. A. Hallmarks of Cancer: The Next Generation. Cell 144, 646–674 (2011).

54. Pernot, S., Evrard, S. & Khatib, A.-M. The Give-and-Take Interaction Between the Tumor Microenvironment and Immune Cells Regulating Tumor Progression and Repression. Front. Immunol. 13, (2022).

55. Gao, Q. et al. Heterotypic CAF-tumor spheroids promote early peritoneal metastatis of ovarian cancer. J Exp Med 216, 688–703 (2019).

56. Islam, M. N. et al. Mitochondrial transfer from bone-marrow–derived stromal cells to pulmonary alveoli protects against acute lung injury. Nat Med 18, 759–765 (2012).

57. Delaunay, S. et al. Mitochondrial RNA modifications shape metabolic plasticity in metastasis. Nature 607, 593–603 (2022).

58. Wang, X. & Gerdes, H.-H. Transfer of mitochondria via tunneling nanotubes rescues apoptotic PC12 cells. Cell Death Differ 22, 1181–1191 (2015).

59. Guan, F. et al. Mitochondrial transfer in tunneling nanotubes—a new target for cancer therapy. J Exp Clin Cancer Res 43, 147 (2024).

60. MacAskill, A. F. & Kittler, J. T. Control of mitochondrial transport and localization in neurons. Trends in Cell Biology 20, 102–112 (2010).

61. Davis, C. O. et al. Transcellular degradation of axonal mitochondria. Proceedings of the National Academy of Sciences 111, 9633–9638 (2014).

62. Lin, R.-Z. et al. Mitochondrial transfer mediates endothelial cell engraftment through mitophagy. Nature 629, 660–668 (2024).

63. Haimovich, G., Dasgupta, S. & Gerst, J. E. RNA transfer through tunneling nanotubes. Biochemical Society Transactions 49, 145–160 (2020).

64. Korenkova, O. et al. Tunneling nanotubes enable intercellular transfer in zebrafish embryos. Dev Cell S1534-5807(24)00635–X (2024) doi:10.1016/j.devcel.2024.10.016.

65. Zheng, K., et al. Long-term Intravital Investigation of an Orthotopic Glioma Mouse Model via Optical Coherence Tomography Angiography. In Vivo 38, 1192–1198 (2024).

66. Schildge, S., Bohrer, C., Beck, K. & Schachtrup, C. Isolation and culture of mouse cortical astrocytes. J Vis Exp 50079 (2013) doi:10.3791/50079.

67. Schindelin, J., et al. Fiji: an open-source platform for biological-image analysis. Nat Methods 9, 676–682 (2012).

68. Ershov, D. et al. TrackMate 7: integrating state-of-the-art segmentation algorithms into tracking pipelines. Nat Methods 19, 829–832 (2022).

69. Sáenz-de-Santa-María, I., Henderson, J. M., Pepe, A. & Zurzolo, C. Identification and Characterization of Tunneling Nanotubes for Intercellular Trafficking. Current Protocols 3, e939 (2023).

70. Cokelaer, T., Desvillechabrol, D., Legendre, R. & Cardon, M. ‘Sequana’: a Set of Snakemake NGS pipelines. Journal of Open Source Software 2, 352 (2017).

71. Köster, J. & Rahmann, S. Snakemake--a scalable bioinformatics workflow engine. Bioinformatics 28, 2520–2522 (2012).

72. Shen, X.-X. et al. Tempo and Mode of Genome Evolution in the Budding Yeast Subphylum. Cell 175, 1533–1545.e20 (2018).

73. Dobin, A. et al. STAR: ultrafast universal RNA-seq aligner. Bioinformatics 29, 15–21 (2013).

74. Love, M. I., Huber, W. & Anders, S. Moderated estimation of fold change and dispersion for RNA-seq data with DESeq2. Genome Biology 15, 550 (2014).

75. Gene Ontology Consortium, T. Expansion of the Gene Ontology knowledgebase and resources. Nucleic Acids Research 45, D331 (2016).

76. Cokelaer, T., Pultz, D., Harder, L. M., Serra-Musach, J. & Saez-Rodriguez, J. BioServices: a common Python package to access biological Web Services programmatically. Bioinformatics 29, 3241–3242 (2013).

